# Efficient approximate Bayesian inference for large, heterogeneous biological datasets

**DOI:** 10.64898/2026.10.05.756786

**Authors:** Jacy T. Zanussi, Christopher E. Miles

**Affiliations:** Center for Complex Biological Systems, University of California, Irvine, CA, USA; Department of Mathematics, University of Utah, Salt Lake City, UT, USA

## Abstract

Simulation-based inference (SBI) offers a route to connect biological data with mechanistic models. At the frontier of biology, datasets continue to grow in scale and richness, and increasingly record heterogeneity between individual observations that a mechanistic model can take into account. Approximate Bayesian Computation (ABC) remains a popular approach to SBI due to its ease of use, but incorporating this heterogeneity often requires a separate simulation of each observation, which is prohibitively expensive at scale. A previously proposed technique, minibatch Sequential Monte Carlo ABC (SMC ABC), reduces this cost by simulating a fixed-size random subset of observations for each proposal, at the price of sampling noise that limits precision. Here, we present an adaptive minibatch approach that uses small batches during early exploration and increases the batch size as the posterior is refined. The adaptive batch size is set by the target precision and does not grow with the size of the dataset, so the savings over full-data SMC ABC increase for larger datasets. We demonstrate the method on synthetic biological models and on real single-cell gene-expression data that vary in scale and type of heterogeneity, including cell geometry and initial condition. In each case, the adaptive scheme obtains posteriors close to those of full-data SMC ABC at a fraction of the cost. Altogether, adaptive minibatching improves the tradeoff between speed and precision in SMC ABC and opens the door to mechanistic inference on large, heterogeneous biological datasets.

## 1 Introduction

Simulation-based inference (SBI) is widely used to connect data with agent-based [1] and other intractable mechanistic models [2, 3]. The ever-growing scale and richness of biological datasets provide opportunities to incorporate more information into increasingly complex models for deeper mechanistic insight. Measurements are now often multimodal, offering additional information about the heterogeneity between individual observations. Heterogeneity in biology is pervasive and often meaningful [4], and mechanistic models increasingly account for it, for example as unobserved cell-to-cell variability in reaction kinetics [5, 6]. When the heterogeneity is measured, it can instead be incorporated directly into the model. For instance, the geometry of an observed cell can define the boundary of a spatial model. However, each observation then needs its own simulation, which makes inference prohibitively expensive for large datasets.

Approximate Bayesian Computation (ABC) is a classical approach to SBI. Its basic premise is to estimate a posterior distribution by repeatedly proposing parameters, simulating data, and retaining proposals whose simulations are sufficiently close to the observations. Although deep-learning-based methods often outperform ABC in direct comparisons [7], ABC remains attractive due to its straightforward implementation and extensive methodological literature. It continues to be used across biology, from systems biology [8, 9] and bioinformatics [10] to epidemiology [11]. A widely used variant, Sequential Monte Carlo ABC (SMC ABC), approximates the posterior efficiently through a sequence of intermediate distributions [12–14]. The particles from one generation guide proposals in the next, while a decreasing acceptance threshold requires progressively closer agreement with the data. Many improvements to ABC and SMC ABC have been proposed, including adapting the summary statistics [15], distance function [16], or proposal kernel [13], and making the sampler [17, 18] or the simulator [19, 20] more efficient. While these efficiency gains can be large, they do not resolve the fundamental computational bottleneck of repeating a separate simulation for each observation of a large dataset.

One way around this bottleneck, ubiquitous in large-scale machine learning, is to subsample the data at each step in a so-called minibatch [21]. Martina Perez, Sailem and Baker [22] proposed simulating a randomly selected batch of observations for each SMC ABC parameter proposal. This reduces the cost of each proposal, but introduces sampling noise into the comparison between simulated and observed data. Because their method uses a fixed batch size throughout inference, it faces an inherent tradeoff between cost and inference quality. Small minibatches are cheap to evaluate, but provide only coarse estimates of the summary statistics to compare with data. Larger minibatches give finer resolution, but save less computation. Subsampling has also been studied in likelihood-based MCMC on large datasets, where the bias it introduces can be analyzed [23, 24] and the subsample can be enlarged adaptively until each accept/reject decision is made with controlled error [25]. In stochastic optimization, a similar approach progressively enlarges the batch when the estimated sampling variance of the gradient becomes large relative to the gradient itself [26]. We take inspiration from these ideas and adapt them to SMC ABC.

We present an adaptive minibatch SMC ABC algorithm that changes the batch size as inference proceeds. Intuitively, coarse selection is sufficient during early exploration, but refining the posterior requires comparisons that represent more of the dataset. We derive a minibatch size update rule that keeps the extra noise from minibatching in balance with the ABC tolerance. Because sampling noise falls with the square root of the batch size, halving the tolerance needs about four times as many observations. Importantly, the cost at a specified precision is therefore independent of the (possibly large) size of the dataset. We test the rule on two synthetic examples and one real-data application. These examples show that the adaptive rule can obtain posterior distributions close to those of the full batch with fewer simulations, resolving the tradeoff between computational cost and inferential precision.

## 2 Adaptive minibatch SMC ABC

### 2.1 Problem setup

We consider a dataset 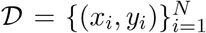 of *N* observations. Each observation pairs a measured outcome *y*_*i*_ with a vector of covariates *x*_*i*_ describing the conditions under which it was made. In the examples below, *x*_*i*_ is the nuclear geometry, including the transcription-site position, or the initial predator and prey counts of a time series, or the time at which a cell was fixed. The model is a stochastic simulator with parameters *θ*. Given *θ* and the covariates *x*_*i*_, it produces an outcome 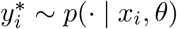. We can draw from *p*(·|*x*_*i*_, *θ*) but cannot evaluate it. The goal is to sample from the posterior

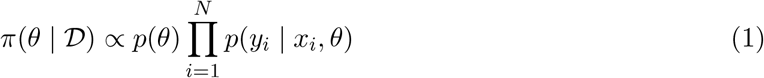

for a prior*p*(*θ*).

### 2.2 ABC, SMC ABC, and minibatching

ABC replaces the likelihood with an acceptance step. A summary function *s*(*y, x*) reduces an observation to a vector of statistics, and *s*_obs_ denotes the summary of the observed data. For a proposed *θ*, we simulate, summarize, and accept the proposal when the simulated summary falls within a tolerance *ϵ* of *s*_obs_. The accepted proposals are draws from

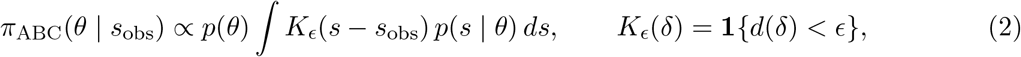

where *d* is a distance on summary differences. The integral is the probability that a simulation at *θ* is accepted. The selection kernel *K*_*ϵ*_ smooths the likelihood on the scale of *ϵ*, so, for summaries that retain the information in the data, the ABC posterior is a broadened version of the posterior in Equation (1). Decreasing *ϵ* sharpens it at the cost of more rejections.

SMC ABC lowers this cost by approaching a small tolerance through a sequence of generations *t* = 0, 1, … [12–14]. The first generation proposes parameters from the prior; later generations perturb the retained particles to propose new parameters. We use a population Monte Carlo sampler with a quantile-based tolerance [16, 27]. With *P* particles and quantile *a*, each generation collects *M* = ⌈*P/a* ⌉proposals that pass the current threshold *ϵ*_*t*_, retains the *P* closest, and uses their largest distance as *ϵ*_*t*+1_. Importance weights correct for drawing from the proposal density *q*_*t*_ rather than the prior. The retained population then guides the next generation. We also resample when the weights become concentrated on too few particles. Additional implementation details are given in Appendix B.4. With *n*_*t*_ = *N*, Algorithm 1 is the standard SMC ABC we build on.

In the setting of Section 2.1, each proposal is evaluated on a minibatch *B* of *n*_*t*_ observations drawn at random, and each selected observation is simulated under its own covariates and compared with its own observed outcome. We average the signed summary differences over the batch before taking the distance (Martina Perez et al. [22] instead average per-observation distances; see Section 4),

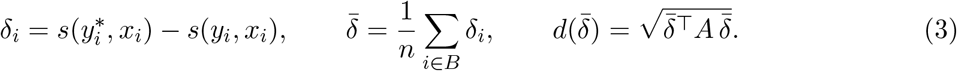

Here *A* is a fixed symmetric positive-semidefinite weight matrix that places the summary components on a common scale. In all our examples, 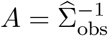 is the inverse of a regularized covariance of the observed summaries, so *d* is a Mahalanobis distance. Diagonal inverse-variance weights are another common choice [16]. We draw *B* uniformly with replacement, which makes the terms of the batch mean independent and identically distributed and simplifies the derivation of the update rule. In practice, this choice seems to have little consequence, and the algorithm works just as well with minibatches chosen without replacement.

### 2.3 Why the batch size should grow

Algorithm 1 with a fixed minibatch size *n*_*t*_ is approximately the approach in [22], aside from some algorithmic differences unrelated to the choice of minibatch. Next, we will motivate why it is beneficial to adapt this size. **Fig**. 1 follows a proposal through one generation: sample *θ*, draw the batch *B*, simulate, compute the distance, accept or reject. Early on, the aim is to separate broadly plausible parameters from poor ones, and a coarse comparison from a few observations is enough. Later generations concentrate on a small region of parameter space, where proposals differ by less than the sampling noise of a small batch. With a constant batch (**Fig**. 1A), the tolerance window narrows across generations while the comparison noise keeps the same width. Proposals are then accepted or rejected by the luck of the batch draw rather than by their fit, and the posterior stops narrowing. Increasing the batch size (**Fig**. 1B) narrows the noise distribution along with the window and lets the finer posterior structure be resolved. A constant batch of the final size would resolve the same structure, but would pay for it from the first generation.

**Fig. 1:**
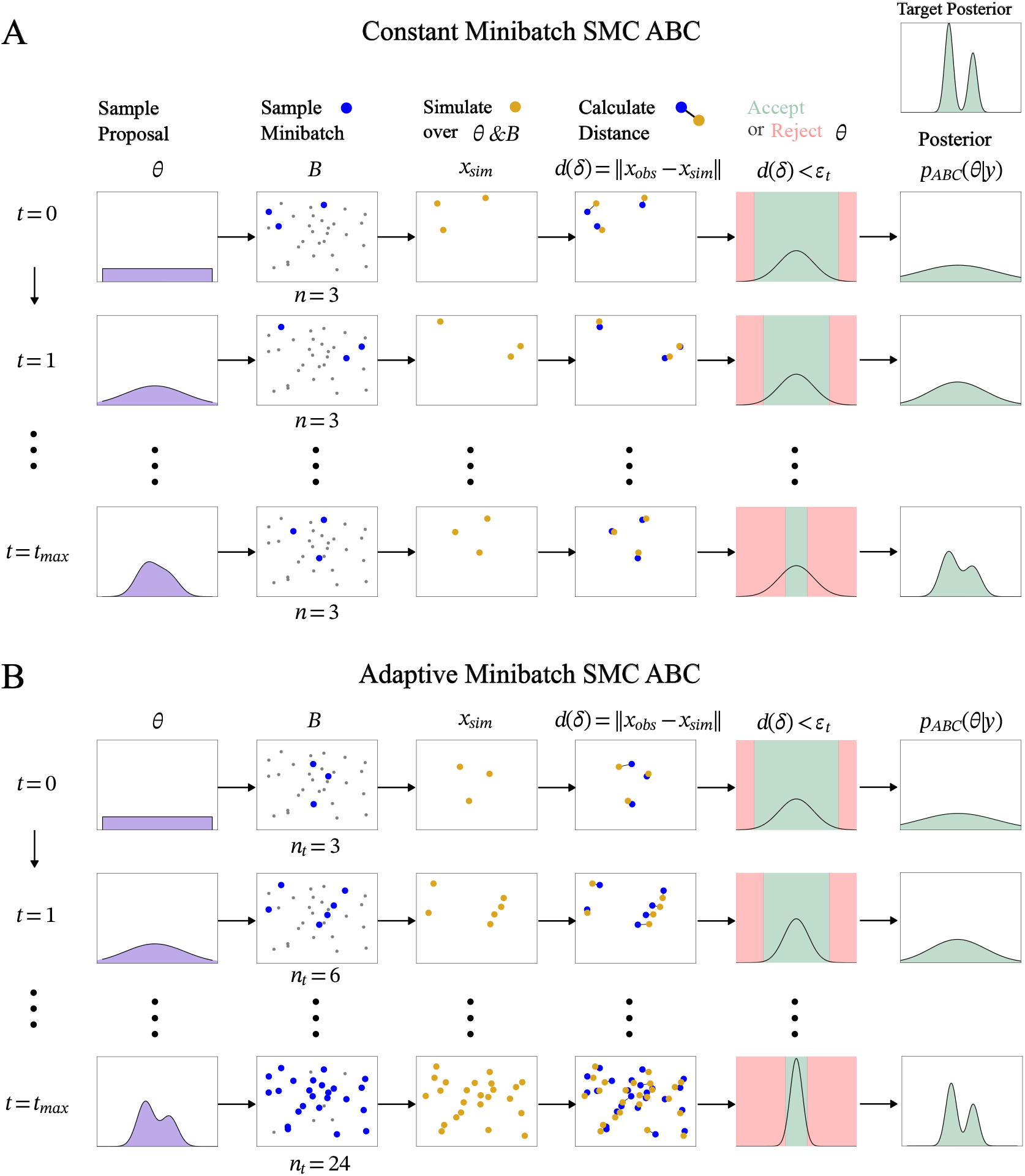
Constant and adaptive minibatch SMC ABC. Each row follows a parameter proposal through batch selection, simulation, distance calculation, and acceptance or rejection. Successive rows show the decreasing tolerance *ϵ*_*t*_ across generations. (A) With a constant batch, the comparison noise (curves in the acceptance column) keeps the same width while the acceptance window (green) narrows, and the posterior stops narrowing. (B) Increasing the batch size narrows the noise with the window, allowing finer posterior structure to be resolved. Posterior sketches are schematic.

#### Algorithm 1

Minibatch SMC ABC

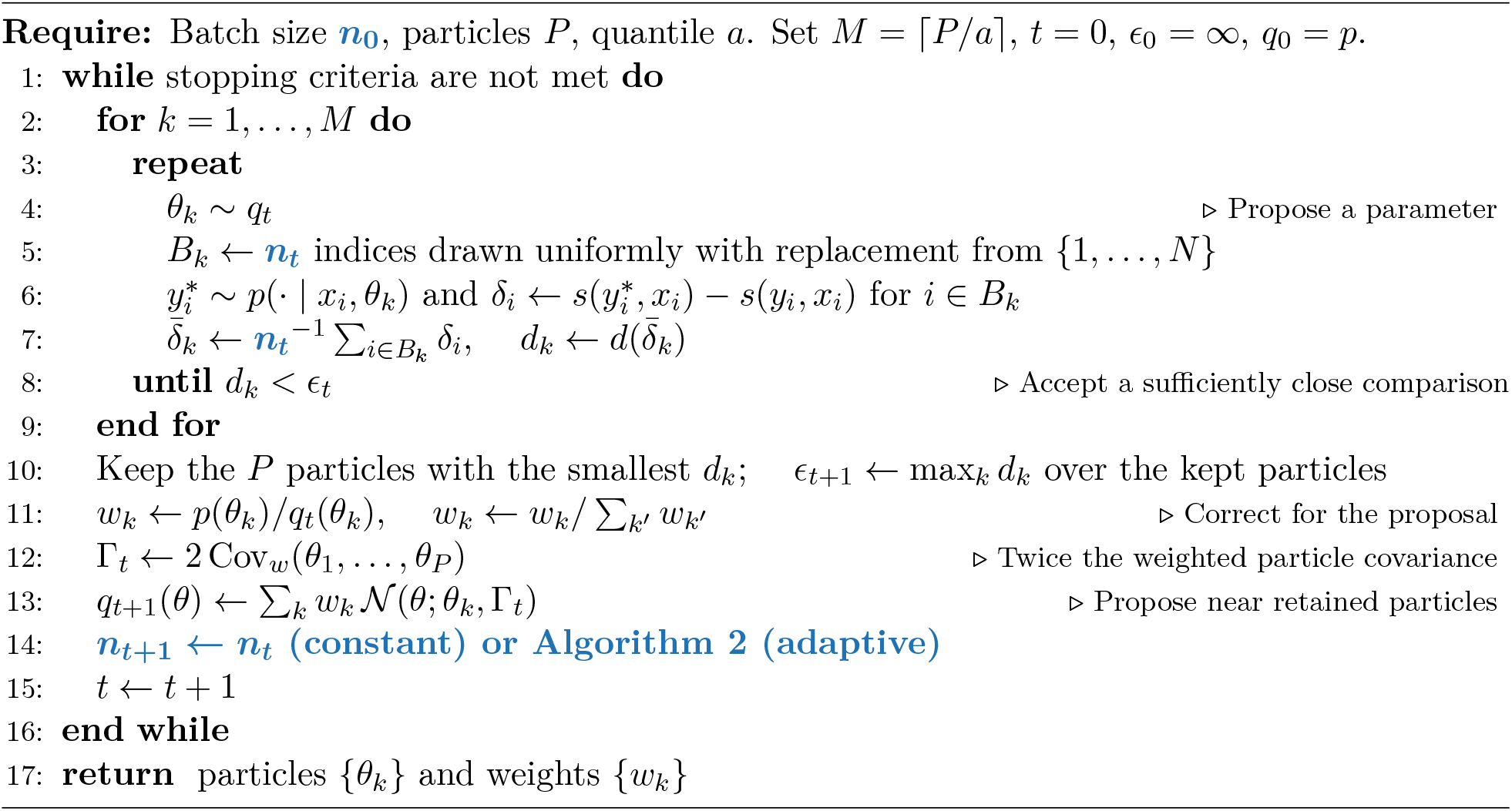

To make this quantitative, we consider the case with a single summary statistic, so *s, s*_obs_, and *ϵ* are scalars. With the hard cutoff, Equation (2) reads

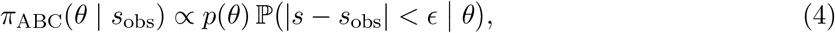

so the acceptance probability determines how each proposal contributes to the posterior. Wilkinson [28] interprets the hard cutoff as adding a uniform error on [−*ϵ, ϵ*]. Minibatching introduces additional sampling noise as well. The general idea of our approach is to control how much this noise smooths the acceptance cutoff.

At a given *θ*, let *µ* = E[*δ*_*i*_] and *σ*^2^ = Var(*δ*_*i*_), where the expectation includes observation sampling and simulator randomness. The central limit theorem gives 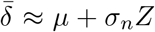, with 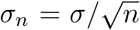 and *Z* ∼N (0, 1). Without sampling noise, the selection kernel is *K*_*ϵ*_(*µ*) = **1** {|*µ*| *< ϵ* }. With sampling noise, the acceptance probability becomes

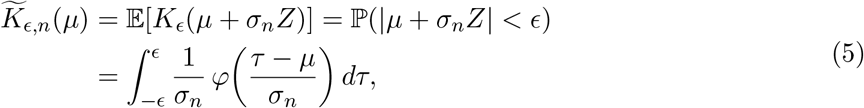

where *φ* is the standard normal density. This is the convolution of the cutoff with the Gaussian sampling density. After dividing by 2*ϵ*, it is the density of the sum of independent uniform tolerance and Gaussian sampling errors. In **Fig**. 1, the acceptance probability is the area under the comparison curve inside the green window. Sampling noise allows proposals with |*µ*| *> ϵ* to be accepted and proposals with |*µ* |*< ϵ* to be rejected.

The amount of smoothing depends on the noise width relative to the tolerance. To see this, write the same acceptance condition in units of *ϵ*:

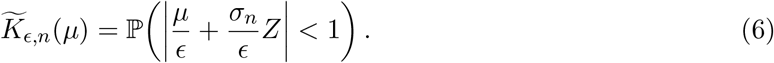

For a given *µ/ϵ*, the acceptance probability depends only on *σ*_*n*_*/ϵ*. With a constant batch and unchanged variance, this ratio grows as the tolerance shrinks (**Fig**. 1A). To preserve the acceptance curve on this scale, we keep the ratio fixed at *c*, so the noise narrows with the window (**Fig**. 1B):

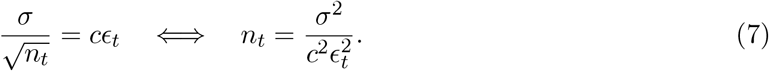

Equivalently, the tolerance-to-noise ratio *r* = *ϵ/σ*_*n*_ stays at 1*/c*. For unchanged variance, halving the tolerance requires four times as many observations.

Even at *n* = *N*, the comparison has sampling variance *σ*^2^*/N*. The full batch is the most precise comparison available and is our reference, so we control only the extra sampling variance introduced by using *n < N, σ*^2^(1*/n* − 1*/N*), and set its standard deviation proportional to the tolerance:

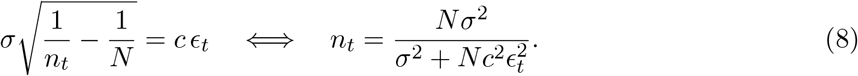

While 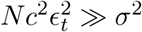, the excess noise dominates the floor, 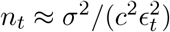, and *r* ≈ 1*/c*: the acceptance curve is approximately preserved on the scale of the tolerance, and the batch size does not depend on *N*. As *ϵ*_*t*_ decreases further, *n*_*t*_ approaches *N* without exceeding it, and the rule hands over to the full batch. Smaller *c* gives larger batches.

For a vector of summaries with weight matrix *A*, the same balance holds with *σ*^2^ replaced by the effective scalar variance

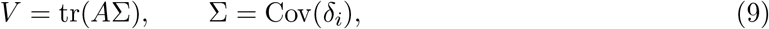

which sums the component variances on the scale used by the distance. Appendix A shows, in the general case, that the batch contributes *V* (1*/n*− 1*/N*) to the expected squared distance in excess of the full batch, and balancing this against 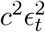 gives

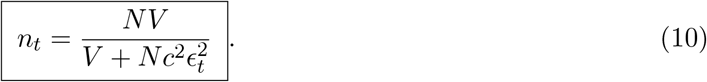

### 2.4 Variance control

The rule needs *V* , which is not known. After generation *t*, we estimate σ from the spread of the differences *δ*_*i*_ within each retained particle’s batch, pooled over particles, and set 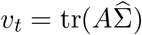 (Appendix A.2). An exponential moving average with weight *λ* stabilizes the estimate across generations,

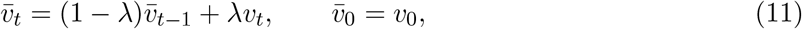

and with *λ* = 1 the current estimate is used without smoothing. The next batch size is then 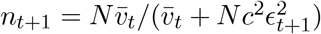, rounded to an integer and restricted to [2, *N*].

The remaining hyperparameters are the initial batch size *n*_0_ and the ratio *c*. Larger *c* gives smaller batches, down to the minimum of two, and smaller *c* gives larger batches, up to *N*. The user can set *c* directly or calibrate it from the first generation by requiring that the first update leave the batch unchanged, *n*_1_ = *n*_0_, which gives

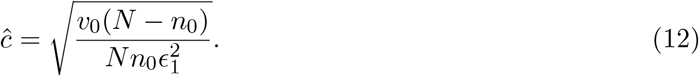

#### Algorithm 2

Variance control update for *n*_*t*+1_

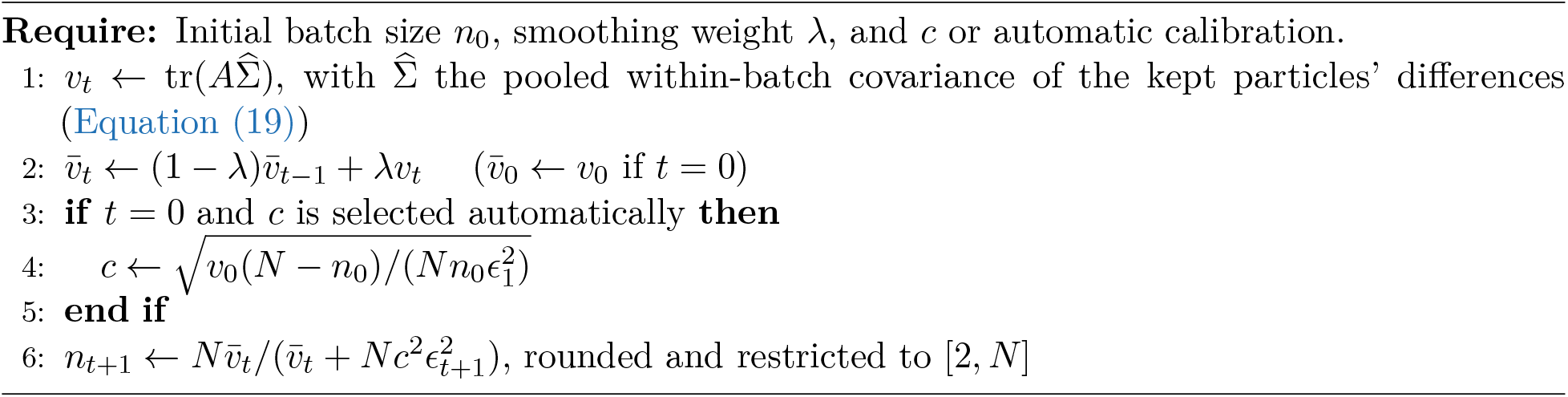

This fixes the ratio of excess noise to tolerance at the value the initial batch reaches at *ϵ*_1_, so the batch grows only as the tolerance tightens further; *c* is then held fixed. Algorithm 2 combines the variance estimate, its smoothing, and the choice of *c* into an automatic update of the batch size after each generation, which we call variance control. The synthetic examples use *λ* = 1, *n*_0_ = 2, and automatic *c* as the default setting. Later, we also vary each hyperparameter in a sweep to show how sensitive the method is to its choice.

## 3 Results

To demonstrate the value of our method in a variety of settings, we compare the adaptive rule with fixed minibatches (including the full batch) on two synthetic models and one real dataset. Throughout, posterior precision is measured by the posterior width *W* = Π _*j*_ *l*_*j*_, the product over the inferred parameters of their marginal 95% highest-density interval lengths *l*_*j*_ (**Fig**. S1). Smaller *W* means tighter marginal distributions, not closer proximity to the true parameters, which are not known in all tests. The computational cost is quantified by wall time or the cumulative number of simulated observations, including those of rejected proposals. Every fit runs to at least generation five and then stops when the acceptance rate falls below 1% or the tolerance falls below the estimated comparison noise. That is, a fit also stops when the signal-to-noise ratio 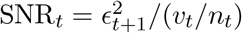, an estimate of *r*^2^ from Section 2.3, falls below one. Additional implementation details can be found in Appendix B.

### 3.1 Transcriptional dynamics with measured geometry

Single-cell measurements resolve the noise in transcription, and fitting mechanistic models of that noise is how the underlying kinetics are inferred [29]. We adapt a model from [30, 31], henceforth referred to as the transcriptional dynamics (TD) model, which describes the spatial pattern of bursty expression of mRNA in cells observed at a single time, as in single-molecule fluorescence in situ hybridization (smFISH) [32]. Importantly, in this example we are inferring dynamics, but the experimental imaging technique fixes the cell, allowing for only one snapshot. We consider a one-dimensional toy version of this model, in which each cell is a nucleus of measured length *L* with a transcription site at measured position *z*. Transcription events occur at *z* with Poisson rate *k*_+_, and each event produces a geometrically distributed number of mRNA with mean *r*_burst_. The mRNA diffuse with diffusivity *D*, degrade at rate *k*_−_, and are exported when they reach either end of the interval (**Fig**. 2). The reactions are

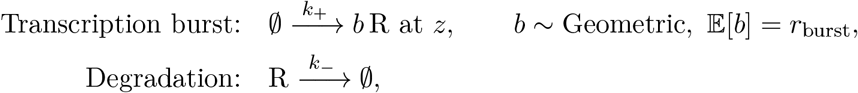

where R is an mRNA molecule. In this example, covariates are *x*_*i*_ = (*L, z*), the geometry of the cell and the transcription-site position, both of which can be resolved from imaging [31]. From snapshots of the surviving mRNA, we infer *k*_+_, *r*_burst_, and *D*, with the degradation rate *k* fixed, assumed to be measured through another experimental modality. The summaries of a cell are its mRNA count, the squared deviation of that count from the mean count of its batch, and the standard deviation of the mRNA positions within the cell (zero for fewer than two mRNA). The first two carry the first and second moments of the counts in a batch, which reflect *k*_+_ and *r*_burst_, and the third reflects *D*. The results of applying our adaptive minibatch method, against the fixed minibatch approach, can be seen in **Fig**. 3. The adaptive rule recovers posterior marginals concentrated around the generating parameters, with widths close to those of the full batch, whereas small fixed batches leave broader distributions. Adaptation achieves this concentration in less wall time (**Fig**. 4A) and with fewer simulated cells (**Fig**. S2). Starting with a small batch therefore preserves much of the computational benefit of minibatching while allowing precise parameter inference later.

**Fig. 2:**
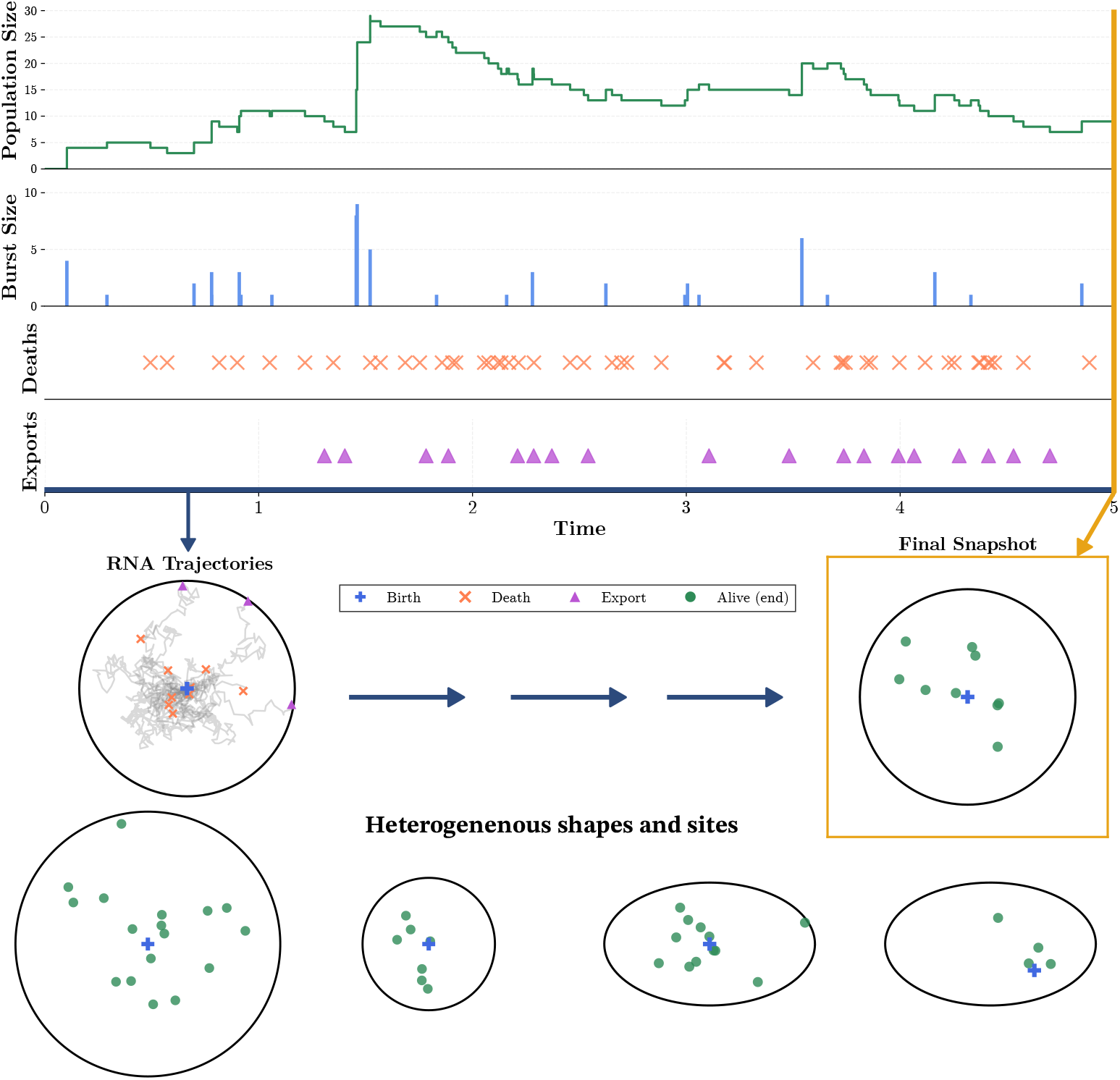
Transcriptional dynamics with measured geometry. (A) Bursty production, diffusion, degradation, and export determine the mRNA present at fixation. (B) Nuclear geometry and transcription-site position affect the observed snapshot. The simulations use a one-dimensional interval with a measured length and site position for each cell.

**Fig. 3:**
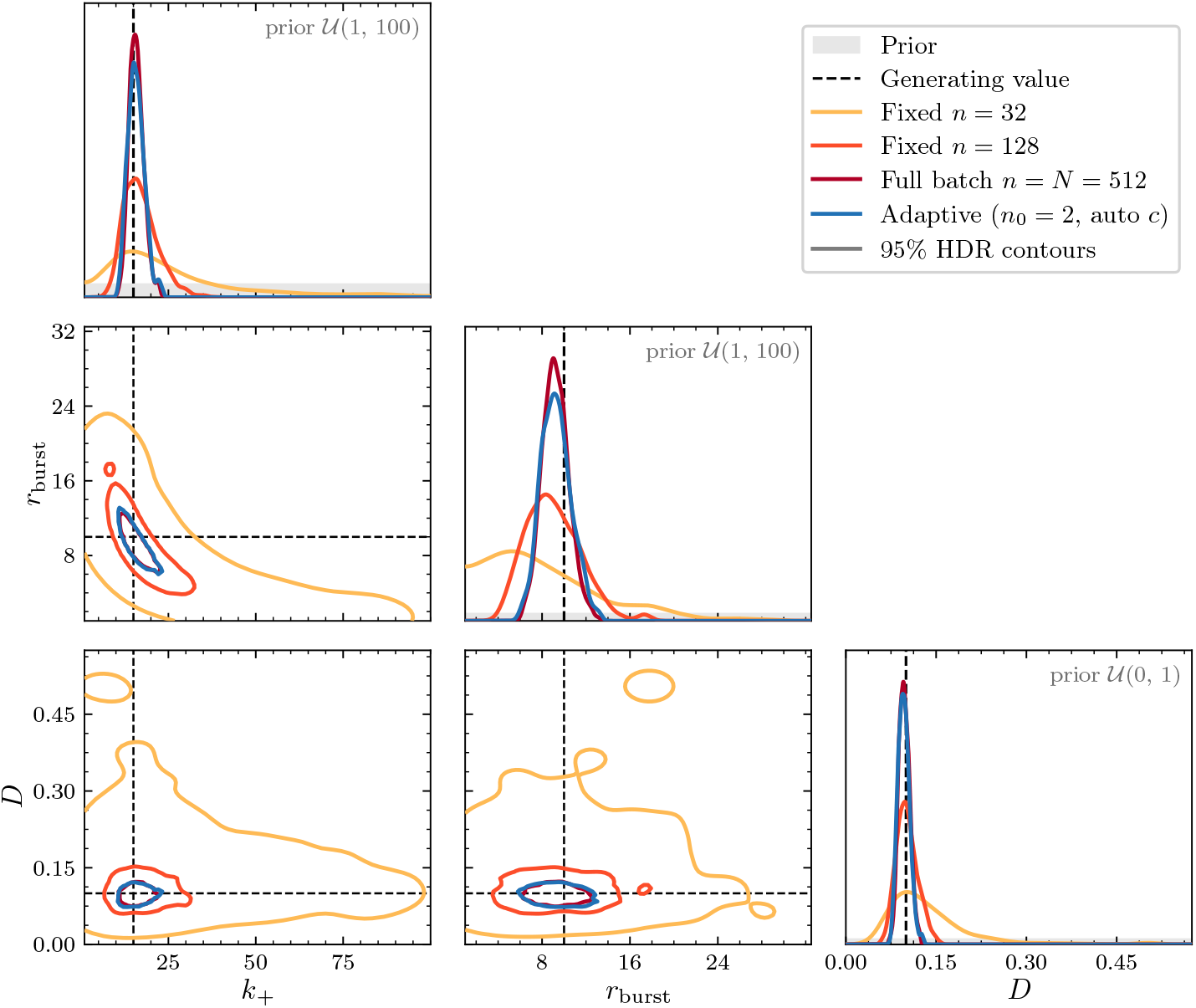
Posterior for the transcriptional dynamics model. The adaptive rule is compared with fixed minibatches and the full batch (*n* = *N* ). The diagonal shows the marginal densities, with grey shading for the prior; the lower panels show, for each pair of parameters, the contour enclosing 95% of each method’s estimated posterior mass. Dashed lines mark the generating parameters. Distributions pool the particles of three seeds at each method’s stopping generation.

**Fig. 4:**
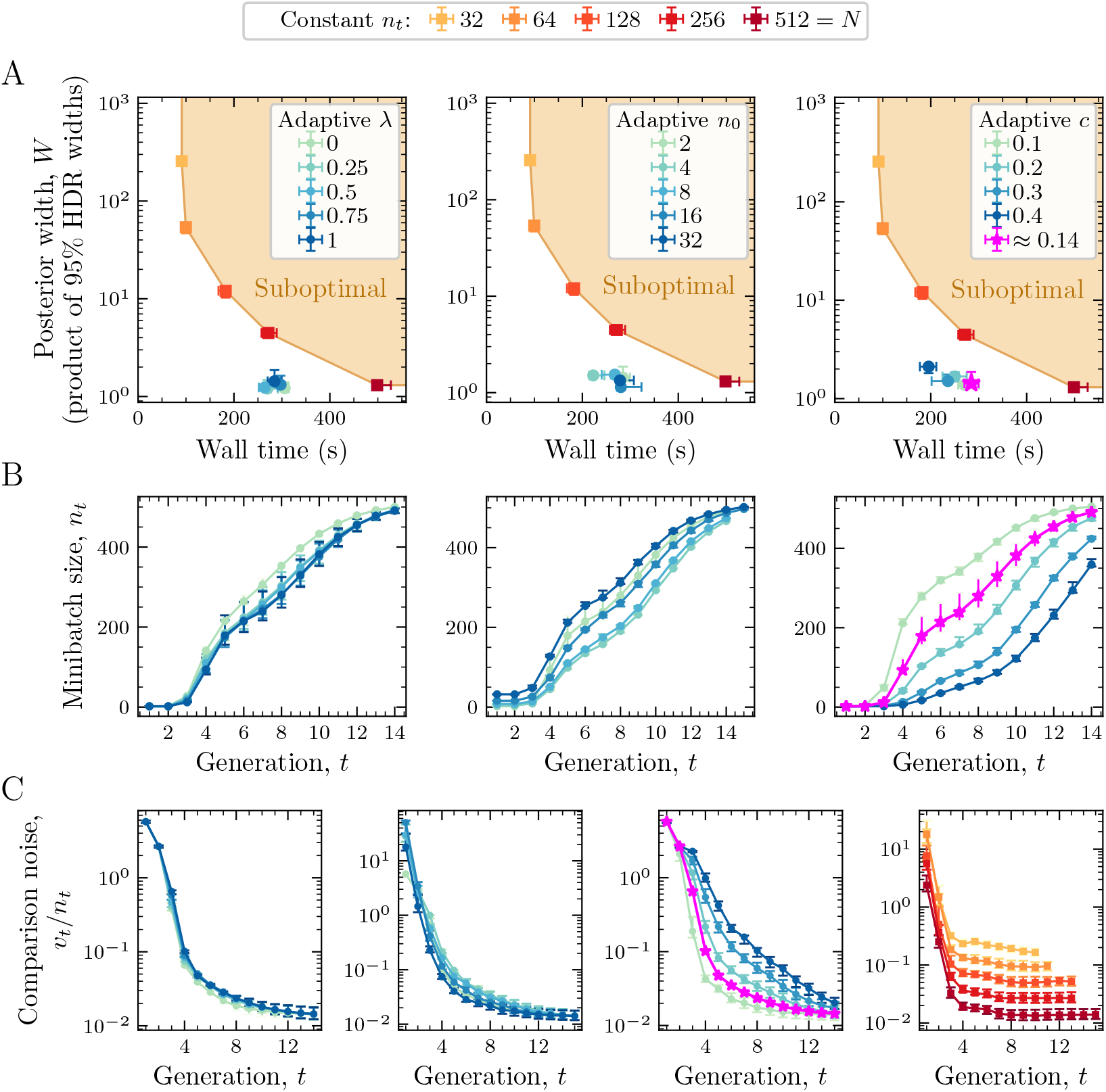
Performance and dynamics of the adaptive rule in the transcriptional dynamics model. (A) Wall time at the stopping generation versus the posterior width *W* , the product of marginal 95% highest-density interval lengths, for sweeps of the smoothing weight *λ*, initial batch size *n*_0_, and noise ratio *c*. Fixed batches *n*_*t*_ provide the comparison; the shaded region is dominated by the fixed-batch frontier. Points are medians over three seeds with quartile bars. (B) Batch size across generations. (C) Estimated comparison noise *v*_*t*_*/n*_*t*_, the minibatch-noise term in the expected squared distance, including a separate fixed-batch panel. Settings and replication are given in Appendix B.

The batch and variance histories explain how this improvement arises (**Fig**. 4B,C). The batch size starts small in the first generations during exploration of the prior. As the tolerance decreases, the adaptive rule increases the batch size and reduces the estimated comparison variance. Small fixed batches retain a higher variance floor, so further tightening of the tolerance yields less posterior shrinkage. These are the same behaviors illustrated in **Fig**. 1.

Across the hyperparameter sweeps the saving is consistent but modest: adaptation reaches the full-batch width in roughly half to two thirds of the wall time, for every choice of *n*_0_, *c*, and *λ* (**Fig**. 4A). We chose the modest size of *N* = 512 cells for the TD dataset so that the full batch is affordable and each comparison can be repeated many times to show the effect of each hyperparameter.

### 3.2 Cost scaling with dataset size

While the excess noise dominates, the adaptive batch size is approximately *V/*(*c*^2^*ϵ*^2^), which does not depend on *N*, so the simulation cost of reaching a given tolerance should be set by the target precision rather than by the dataset. This feature of our approach should allow far larger efficiency gains on large datasets than the modest gains seen in the hyperparameter sweeps. We test this with TD datasets of *N* = 512 to 32,768 cells, comparing the full batch, a fixed batch of *n* = 512, and the adaptive rule (**Fig**. 5). For each fit we record the cumulative simulated cells at the first generation whose posterior width *W* falls below a target. The full-batch cost grows in proportion to *N* at every target. The adaptive cost grows much more slowly than *N*, by a factor of two to three over the 64-fold range at the looser targets, giving a full-batch-to-adaptive cost ratio that rises from below two at *N* = 512 to between six and sixty at *N* = 32,768. Wall time follows the same pattern (**Fig**. 5B): the adaptive rule reaches each target in about half the full-batch time at *N* = 512 and in one tenth to one eightieth of it at *N* = 32,768. The fixed batch is as cheap as the adaptive rule at loose targets but stalls at a width near 1.5 at every *N*, and never reaches the two tighter targets, whereas the adaptive rule reaches the same targets as the full batch, although at *N* = 8,192 one of its three seeds stops just above the tightest. Altogether, the adaptive rule keeps the low early cost of a small batch without its precision floor, and its cost at a given precision grows only slowly with *N*.

**Fig. 5:**
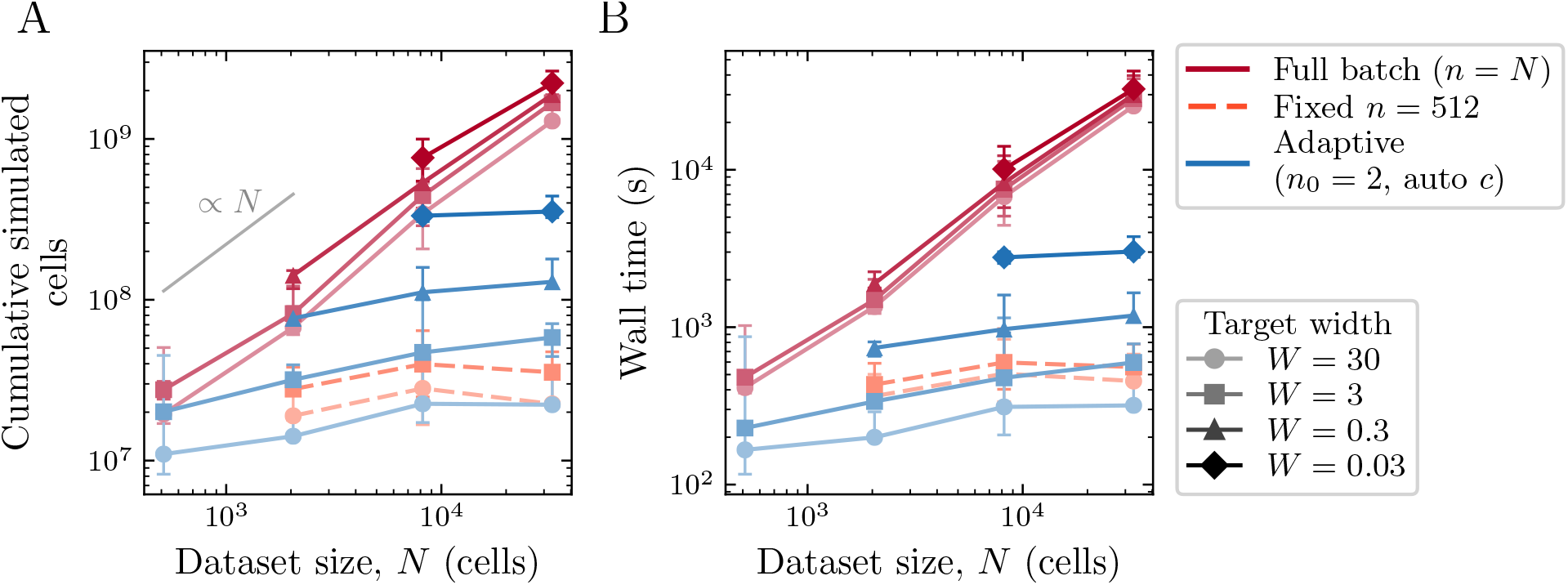
Simulation cost against dataset size in the transcriptional dynamics model. (A) Cumulative simulated cells at the first generation reaching a target posterior width *W* (product of marginal 95% highest-density interval lengths) for the full batch, a fixed batch of *n* = 512, and the adaptive rule, on datasets of *N* = 512 to 32,768 cells. Points are medians, with range bars, over the seeds that reached the target, out of three; a curve ends where no seed reaches the target. The grey segment shows growth proportional to *N*. (B) Wall time at the same generation.

### 3.3 Robustness to model kinetics and heterogeneity

The comparisons so far use one generating parameter set. To ask how robust the result is, we test large perturbations to the model kinetics (**Fig**. 6) and changes in the distribution of nuclear geometries (**Fig**. S3). The adaptive rule improves the cost–shrinkage tradeoff in nearly every kinetic setting. The size of the gain varies between settings without an obvious pattern in the kinetics, and at the lowest diffusivity the methods perform similarly. Changing the shape and scale of the nuclear-length distribution, and with them the mean and spread of cell sizes, gives the same picture (**Fig**. S3): in each of the nine combinations, adaptation reaches widths near the full-batch width in less wall time, although with *n*_0_ = 2 it stops early in one of them. Adaptation is therefore useful across a range of kinetics and geometries, without requiring a batch schedule tuned to one parameter set.

**Fig. 6:**
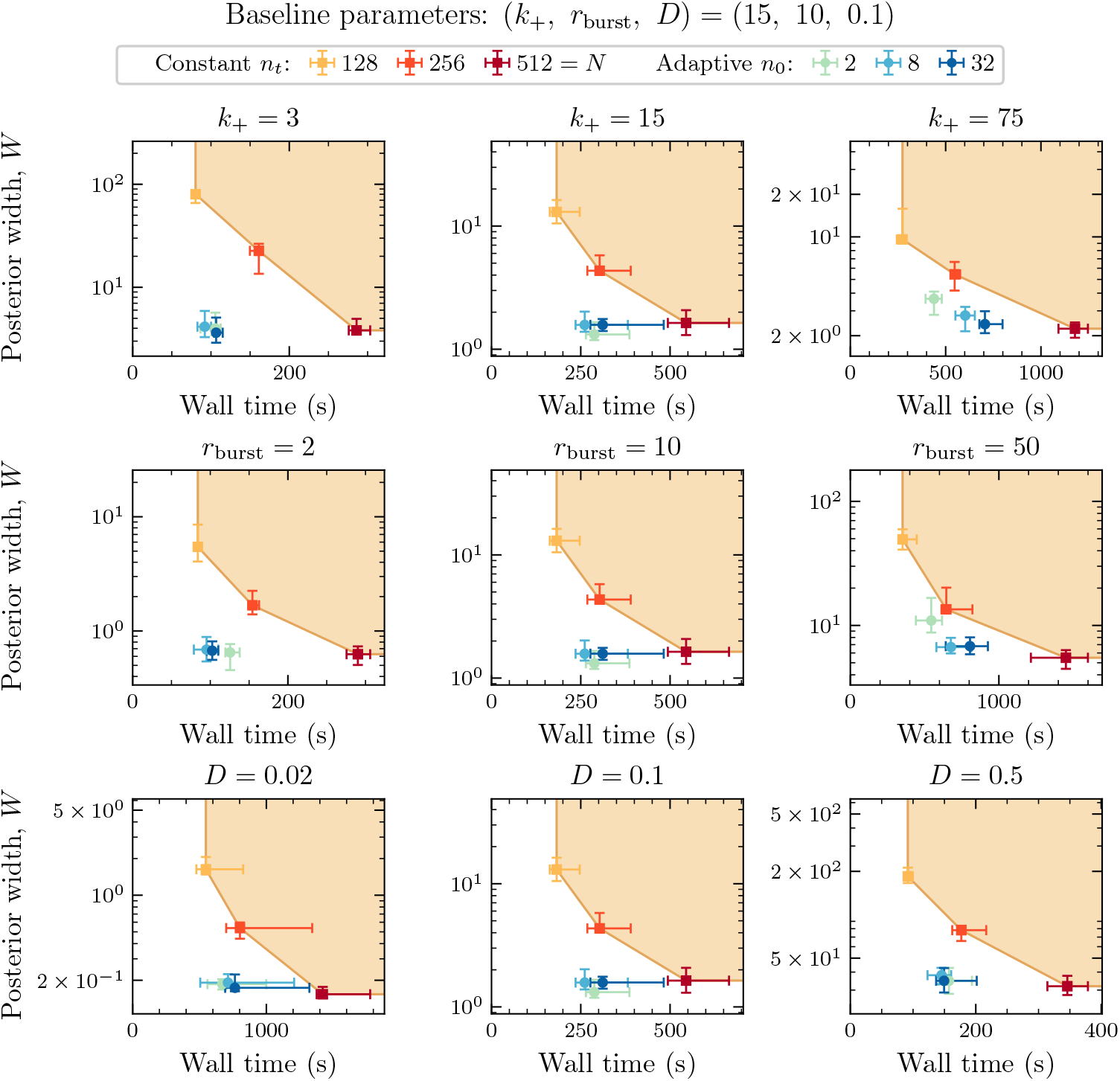
Robustness to changes in transcriptional kinetics. Each panel varies one generating parameter (*k*_+_, *r*_burst_, or *D*) with the others at their baseline values, and shows wall time at the stopping generation against the posterior width *W* for fixed batches *n*_*t*_ and the adaptive rule with initial batch *n*_0_ and automatic *c*. Points are medians over ten paired data/SMC seeds with quartile bars; the shaded region is dominated by the fixed-batch frontier.

### 3.4 Stochastic Lotka–Volterra with measured initial conditions

The stochastic Lotka–Volterra (LV) model provides a second setting, in which measured initial conditions generate heterogeneous trajectories. Here we consider predator–prey time series, each starting from its own observed pair of population counts (**Fig**. 7), similar to the scratch assays of Martina Perez et al. [22], whose simulations begin from the observed initial cell distribution of each well.

**Fig. 7:**
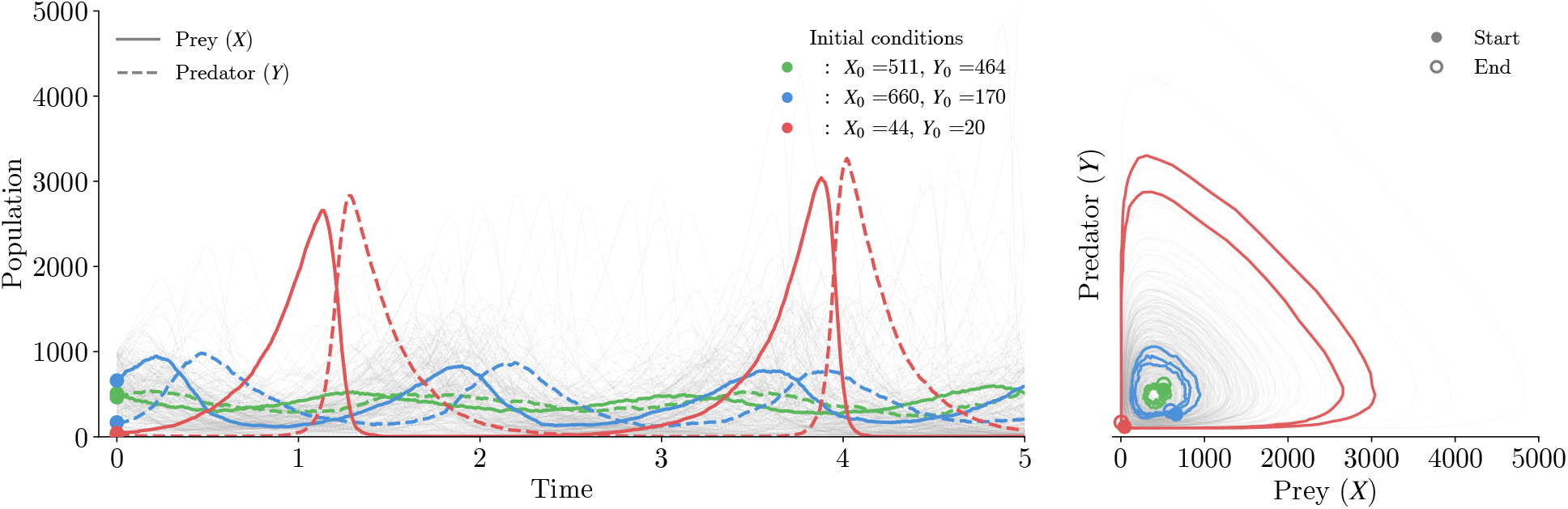
Stochastic Lotka–Volterra dynamics with heterogeneous initial conditions. Different initial populations produce different trajectories. Each trajectory contributes a summary vector, and its measured initial prey and predator counts initialize the corresponding simulation.

In this model, we track two interacting populations over time. Each trajectory follows prey *X* and predators *Y* through the reactions

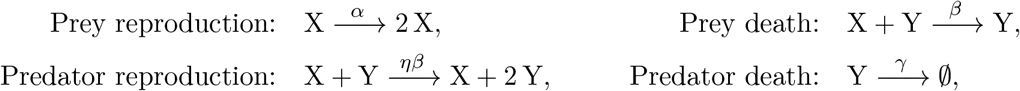

simulated by tau-leaping [33]. We prevent extinction by capping the number of deaths in each time step so that neither population falls below one. The covariates are the initial counts *x*_*i*_ = (*X*_0_, *Y*_0_), and different initial conditions produce very different dynamics, with some populations spiking and hovering near extinction and others staying nearly flat. We infer the prey birth rate *α*, predation rate *β*, and predator death rate *γ*, with the conversion factor *η* fixed. The seven summaries of a trajectory are the log mean and log variance of each population, the autocorrelation of each population at lag 0.1, and the prey–predator cross-correlation, computed up to the first sampled time at which either population reaches one individual.

The adaptive rule again produces posterior marginals concentrated around the generating parameters and close to those of the full batch, while small fixed batches remain broader (**Fig**. 8).

**Fig. 8:**
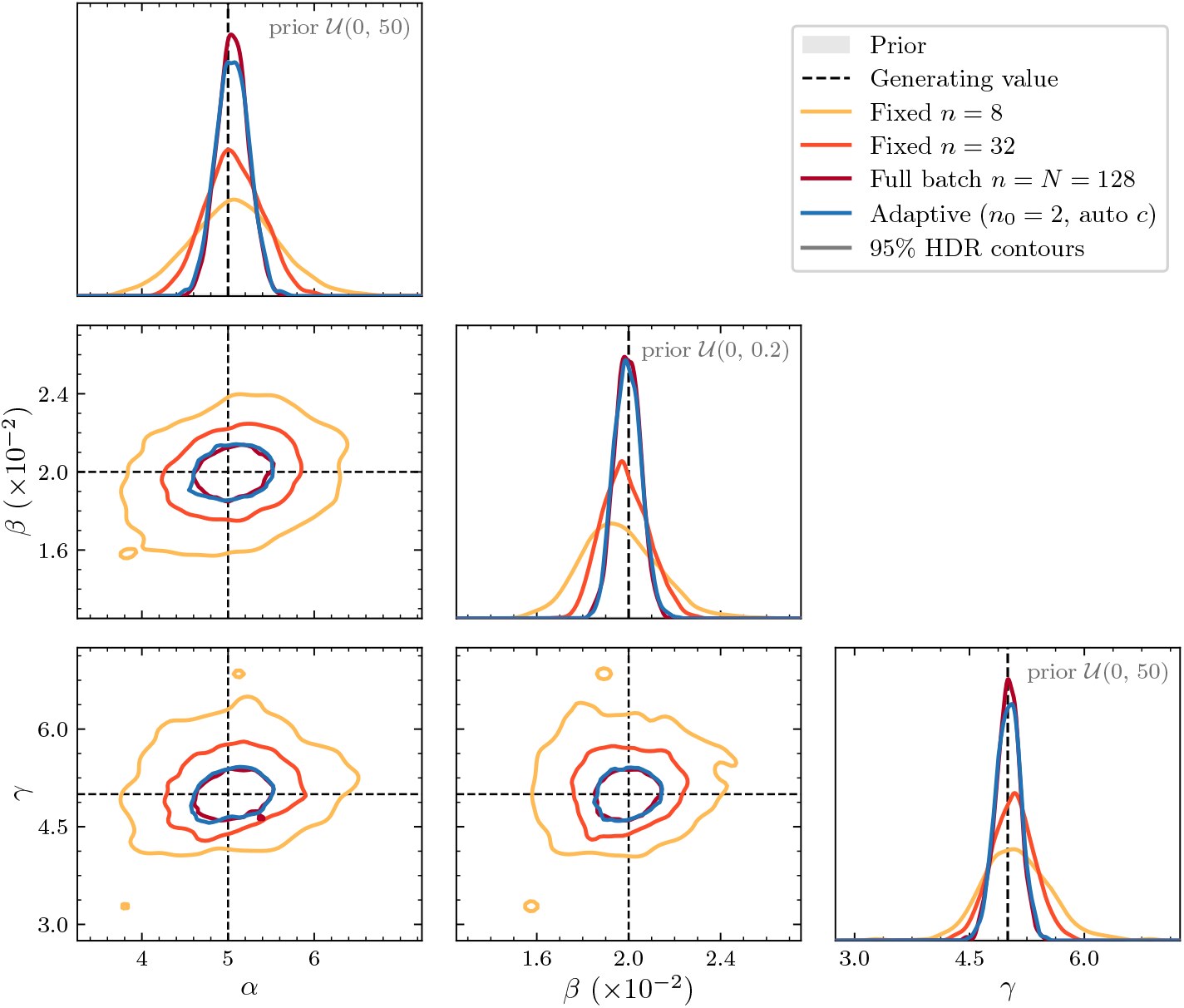
Posterior for the stochastic Lotka–Volterra model, as in **Fig**. 3: marginal densities on the diagonal and 95% contours for each pair of parameters, pooling the particles of three seeds at each method’s stopping generation. Dashed lines mark the generating parameters. The uniform priors are too flat to be visible at this scale; their ranges are given in each diagonal panel.

The cost comparison shows that adaptation reaches narrow posteriors in less wall time (**Fig**. 9A) and with fewer simulated trajectories (**Fig**. S4).

**Fig. 9:**
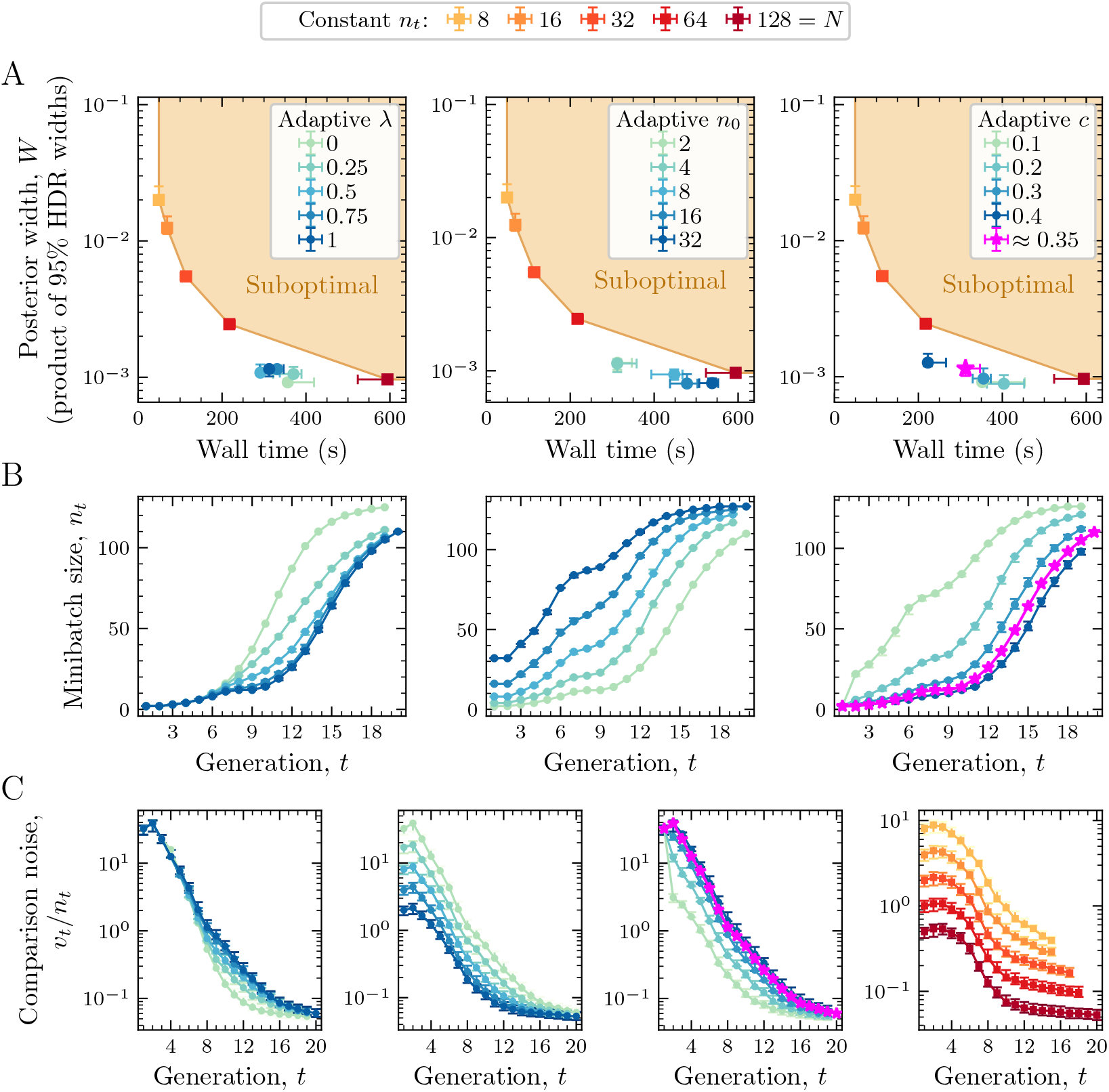
Performance and dynamics of the adaptive rule in the stochastic Lotka–Volterra model. (A) Wall time at the stopping generation versus the posterior width *W* for sweeps of *λ, n*_0_, and *c*, with fixed batches for comparison. (B) Batch size across generations. (C) Estimated comparison noise *v*_*t*_*/n*_*t*_, including a separate fixed-batch panel, as in **Fig**. 4.

We observe much of the same behavior as in the TD model. The adaptive rule increases the batch size as the tolerance decreases, bringing the comparison variance toward the full-batch level (**Fig**. 9B,C). The initial batch size and noise ratio have a more noticeable effect on performance in this example. Changes to the kinetic parameters and to the range of initial populations give the same picture (**Figs**. S5 and S6).

### 3.5 Real single-cell *lacZ* transcription snapshots

In our last demonstration, we apply our method to a previously analyzed dataset of slow-growth *E. coli lacZ* snapshots measured by Wang et al. [34], using the source data distributed with Kilic et al. [35]. In this dataset, the covariate is the fixation time *x*_*i*_ = *t*_*i*_, and a selected cell is simulated up to its own *t*_*i*_ (**Fig**. 10A). The model is a standard telegraph model [36] for transcriptional dynamics, where the promoter switches between two states with different mRNA production rates. The promoter *G* and mature mRNA R follow the reactions

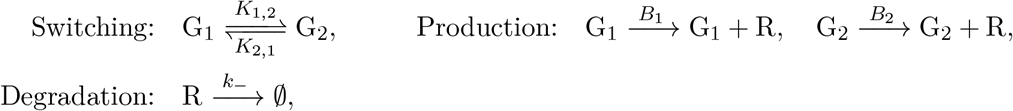

**Fig. 10:**
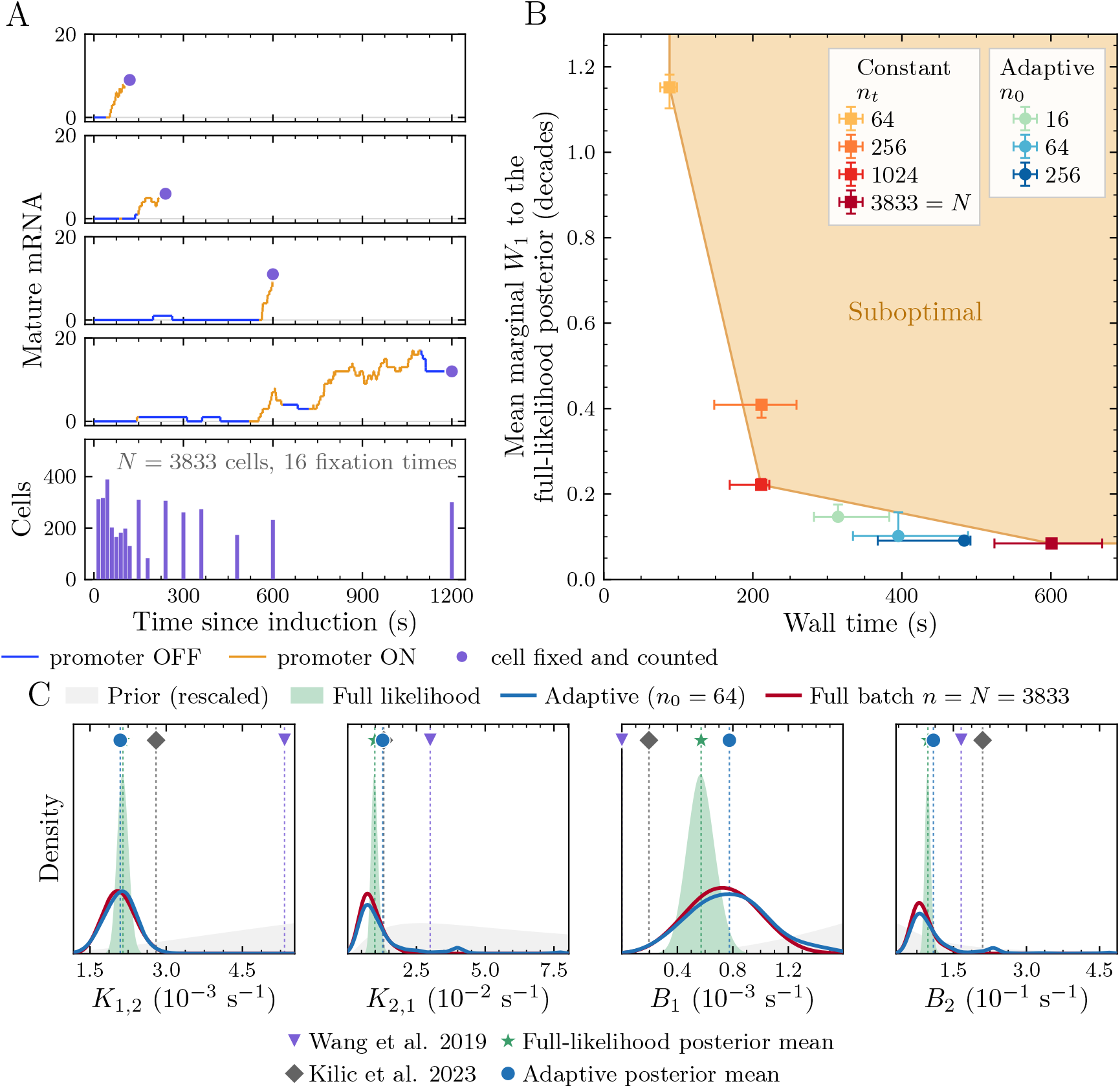
The adaptive rule on real single-cell *lacZ* snapshots. (A) Top: simulated cells run only until their own fixation time, where the mature mRNA count is recorded; the trace is colored by the promoter state. Bottom: the observed dataset, *N* = 3833 cells spread over 16 fixation times, so every observation carries its own simulation length. (B) Wall time at the stopping generation versus mean marginal Wasserstein distance to the likelihood-based posterior, measured in log_10_ rate coordinates. Points show medians over five runs and bars their range; the shaded region is dominated by the fixed-batch frontier. (C) Kernel-smoothed rate marginals pooled across the five runs, on a linear rate scale, with *n*_0_ = 64 for the adaptive rule. The green fill shows the likelihood reference and the green star its mean; the light grey fill shows the shape of the prior, rescaled to be visible. Markers above the densities, with dotted guide lines, give the estimates of Wang et al. [34], the posterior means of the Fig. 4 chain of Kilic et al. [35], the likelihood-based posterior mean, and the adaptive posterior mean. Literature estimates provide context under different modeling assumptions, as detailed in Appendix B.3.

simulated exactly by the Gillespie algorithm [37] from state *G*_1_ with no mRNA at induction. We infer the switching rates *K*_1,2_ and *K*_2,1_ and production rates *B*_1_ and *B*_2_, all in s^−1^, with the degradation rate *k* _−_fixed. The summary of a cell is the vector of indicators **1 {**count≤*m}* for *m* = 0, 1, 3, 7, 15, placed in the block of its fixation time, so that the batch mean is the empirical count distribution at each time; with 16 fixation times this gives 80 summaries.

Real data have no generating parameters to recover. For this model, however, the chemical master equation gives the likelihood, which we evaluate on a truncated state space in the manner of finite state projection [38], as Kilic et al. [35] also do, so we can sample the likelihood-based posterior by MCMC and use it as the reference. The reference separates two errors: the error of ABC itself, and the error that adaptation adds within ABC. We measure the accuracy of an ABC posterior by the 1-Wasserstein distance between its marginal and the reference marginal for each rate, averaged over the four rates, with *W*_1_ = ∫ |*F*_ABC_(*u*) − *F*_ref_(*u*) | *du*, where *F*_ABC_ and *F*_ref_ are the cumulative distribution functions of *u* under the ABC and reference posteriors. This distance to the likelihood posterior is the closest available substitute for distance to the truth, and it compares every method against the same target. Adaptive and full-batch ABC produce closely agreeing posterior marginals, in the same region as the likelihood reference (**Fig**. 10C). Both ABC distributions are broader than the likelihood reference, as expected from the ABC approximation, but their posterior means lie close to the likelihood mean, with the largest offset in *B*_1_. The close agreement between the two ABC posteriors indicates that adaptation introduces little additional discrepancy at the precision reached here.

The adaptive rule reaches this posterior at lower cost (**Fig**. 10B). With *n*_0_ = 64, it stops after about 34% less wall time than the full batch, having simulated about 33% fewer cells, with a similar distance to the likelihood reference. The gain is larger earlier in inference: at intermediate distances, the adaptive rule uses roughly a third to a half as many simulated cells as the full batch (**Fig**. S7). The point estimates tell the same story (**Fig**. S8): the adaptive posterior means settle at the full-batch values after about half as many simulated cells, and reach the likelihood mean of *K*_1,2_ after less than half as many, while small fixed batches, though inexpensive, stop far from it in every rate. This example shows both uses of adaptation: obtaining a moderately precise posterior cheaply and continuing toward full-batch precision when the application requires it.

## 4 Discussion

In this paper, we introduced a rule for adaptively adjusting the minibatch size at each generation of SMC ABC. Larger biological datasets offer more information for fitting mechanistic models, but when each observation must be simulated under its own measured conditions, reproducing the dataset for every proposal is expensive. The adaptive rule uses small batches while the posterior is broad and increases the batch size as the tolerance tightens. In the three examples it obtains distributions close to the full-batch result with fewer simulations, while small fixed batches stall at a precision floor. The batch schedule is set by the algorithm rather than tuned by hand, and while the excess noise dominates, the batch size *V/*(*c*^2^*ϵ*^2^) does not depend on *N*. The cost of a given precision therefore grows only slowly with the size of the dataset, and the savings over the full batch grow with it. Adaptive minibatching thus brings precise mechanistic inference within reach for large datasets in which every observation carries its own measured heterogeneity, a setting where simulating the full dataset for each proposal is prohibitive.

One important caveat of our approach is that the saving depends on the precision target. Once the tolerance is small enough that *Nc*^2^*ϵ*^2^ is comparable to *V* , the batch approaches *N* and the late generations dominate the total cost. The saving to full-batch precision is then more modest than the saving to an intermediate precision, as the *E. coli* example shows within a single inference problem. Which precision is needed should be decided by the scientific question. The authors of [22] chose a fixed batch size after finding that posterior means and variances changed little with further increases, chose the number of generations from the change between successive posteriors, and showed that minibatching retained useful mechanistic differences between experimental conditions. Our method lets the required precision set the simulation effort during the run instead.

Biological heterogeneity can be modeled in many ways, and the choice shapes both the inference and its cost. We have treated heterogeneity that is measured and enters the simulator as a covariate, and compared simulations to observations one to one. We average the signed summary differences over the batch before taking the distance, ∥*n*^−1^ Σ_*j*_ *δ*_*j*_ ∥^2^. Martina Perez et al. [22] instead average per-observation discrepancies, *n*^−1^ Σ_*j*_ ∥*δ*_*j*_∥ ^2^. The latter retains individual mismatches that may cancel in the former, but also carries the variation between stochastic realizations, which does not average out with more observations. In our design the minibatch noise is a bias that shrinks with the batch, so the full batch is the natural endpoint; in theirs it is a fluctuation around a floor that the batch size does not lower, and an adaptive rule would have to work relative to that floor. Heterogeneity can also be the object of inference rather than an input. For instance, subpopulations with different kinetics, such as cell types with different RNA dynamics, call for a distribution of cell-specific rates in the model [5], and hierarchical models estimate the variation between experimental replicates from snapshot data [39]. We have not explored how such models interact with adaptive minibatching. Measurement noise is also notably absent from our simulators, but it can be built into ABC [8] and would enter the comparison variance like any other source of simulator noise.

Adaptive minibatching is simple to add to SMC ABC and uses quantities already available from the simulated–observed comparisons. A small batch can be seen as a low-fidelity version of the full-data comparison, with the batch size playing the role of fidelity. Multilevel and multifidelity schemes combine cheap, approximate simulations with fewer exact ones [19, 40], and could be adapted to mix batch sizes within a generation, for example by simulating the full batch only for proposals that look promising on a small batch. The question of how much simulation to spend at each stage also arises in sequential neural inference [41, 42] and in methods that learn the posterior directly from simulated summary statistics [43], where the same allocation may carry the computational benefit beyond the ABC setting studied here.

## Code and data availability

Code is available at https://github.com/JacyZanussi/adaptive_minibatch_smc_abc. The experimental data are from Wang et al. [34], distributed with the source data and code of Kilic et al. [35].

## Acknowledgments

JTZ was supported by the NIH NIGMS training grant T32GM136624. CEM was supported by NSF CAREER DMS-2545859 and NSF/NIGMS DMS-2451263. We thank Dhara Hansalia, Christopher Au, and Luca Valdevit for preliminary work on this project as part of the 2024 Math ExpLR program at UC Irvine. This work utilized the infrastructure for high-performance and high-throughput computing, research data storage and analysis, and scientific software tool integration built, operated, and updated by the Research Cyberinfrastructure Center (RCIC) at the University of California, Irvine (UCI).

## A Derivation of the variance control rule

### A.1 Excess variance of the minibatch comparison

We fix the dataset *D*, a proposal *θ*, and the weight matrix *A*, and take all expectations over the batch draw and the simulator. Each term *δ*_*i*_ of the batch mean in Equation (3) selects an observation *i* uniformly from {1, … , *N}* and simulates it once, 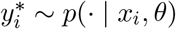. For a fixed observation *i*, the mean and covariance of *δ*_*i*_ over the simulator are

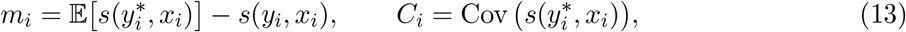

and 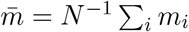 is the average of the means over observations. Over the random choice of *i* as well, the law of total covariance gives

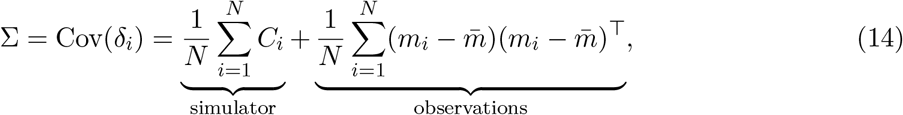

so σ collects the simulator noise at each observation and the variation of the expected difference across observations. Because the batch is drawn with replacement and each draw is simulated independently, the *n* terms of 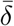 are independent and identically distributed, and the batch mean has

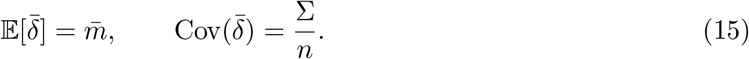

The expected squared distance follows from 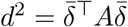:

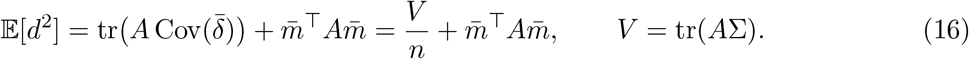

The mismatch term 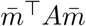 does not depend on the batch size, so the batch of size *n* contributes

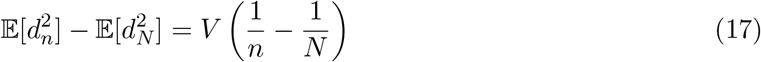

to the expected squared distance in excess of the full batch. Setting this excess equal to 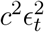 at generation *t* gives

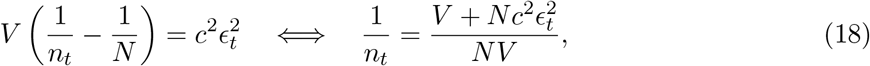

and taking the reciprocal yields Equation (10). For a single standardized summary, *V* = *σ*^2^ and this is Equation (8).

### A.2 Estimating *V*

Let *δ*_*ki*_, *i* ∈*B*_*k*_, be the differences stored for retained particle *k* and 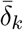 their mean. Centering each batch separately removes the mismatch 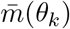, which differs between particles, and pooling gives

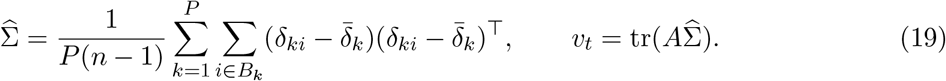

Equation (17) holds at a fixed parameter before acceptance selection; the estimate pools over the accepted particles of the generation, which is what the algorithm has available.

## B Model and algorithm settings

### B.1 Transcriptional dynamics

We simulate each cell from an empty nucleus to the imaging time *T* = 5. The burst times, burst sizes, and mRNA lifetimes are sampled exactly. Each mRNA then diffuses from *z* by Brownian steps with time step Δ*t* = 0.01 and is exported when it first leaves (0, *L*). The data are the positions of the mRNA that have survived degradation and export at time *T* . We infer (*k*_+_, *r*_burst_, *D*) over uniform priors on [1, 100] × [1, 100] × [0, 1] and fix the degradation rate at *k* _−_= 5. The heterogeneity is in nuclear size and transcription-site location: we draw the length *L* from a Gamma(10, 0.1) distribution (shape and scale) and set the site at *z* = *LU* with *U* ∼Beta(2, 2). The synthetic dataset has *N* = 512 cells generated at (*k*_+_, *r*_burst_, *D*) = (15, 10, 0.1).

### B.2 Stochastic Lotka–Volterra

We simulate each trajectory from its initial counts to the final time *T* = 5 with tau-leaping time step Δ*t* = 0.01. In each step, the numbers of prey and predator deaths are capped so that neither population falls below one. We infer (*α, β, γ*) over uniform priors on [0, 50] × [0, 0.2] × [0, 50] and fix the conversion factor at *η* = 1. The heterogeneity is in the initial condition: we draw (*X*_0_, *Y*_0_) independently and uniformly from the integers 1 to 1000. The synthetic dataset has *N* = 128 trajectories generated at (*α, β, γ*) = (5, 0.02, 5).

### B.3 Telegraph model for *lacZ* transcription

We follow the prior of Kilic et al. [35]: the four rates have independent *N* (−1, 0.5^2^) priors on their log_10_ values in s^−1^. We fix *k*_−_ = 0.00533 s^−1^, the slow-growth degradation rate (0.32 min^−1^) of the two-state fit of Wang et al. [34]. The initial state, *G*_1_ with no mRNA, matches the cells imaged at induction, none of which contains mRNA. The data are the mature mRNA counts of *N* = 3833 cells at 16 fixation times from 15 to 1200 s.

The likelihood reference uses the same prior, initial state, and *k*_−_, with the state space truncated at 140 mRNA. We sample it by adaptive Metropolis [44] in log_10_ coordinates, running four chains of 40,000 iterations, discarding the first 15,000, and keeping every fourth draw, for 25,000 draws in total 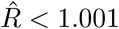 for every rate).

The literature markers come from analyses that differ slightly from ours. Wang et al. [34] fit a model that also includes an induction delay, finite elongation, RNA retention and permanently inactive cells. The Kilic et al. [35] markers are posterior means of the MCMC chain distributed as source data for their Fig. 4, which analyzes the same counts in our parameter order, averaged over the two-state samples after a 30% burn-in, with the state of lower production rate taken as *G*_1_. That chain also samples the number of promoter states and the degradation rate, which settles near 0.015 s^−1^, and their released code starts cells with some mRNA present. We therefore treat the literature markers as context and emphasize that the full likelihood reference is the most appropriate comparison for our ABC posteriors.

### B.4 Algorithmic settings

**Tab**. 1 lists the settings for each model. All fits use the quantile *a* = 0.5, the Mahalanobis weight 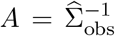 with 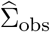 a regularized covariance of the observed summaries, and the perturbation kernel of Algorithm 1, with resampling as described below. In the stopping rule of Section 3, the acceptance rate is *M/R*_*t*_, where *R*_*t*_ is the number of proposals simulated. The robustness sweeps also cap fits at 25 generations and six hours; no fit reached either cap. Wall times are measured on a compute cluster with eight worker processes per fit.

**Table 1.** Settings for the three examples. Lists give the values swept one at a time.

|  | Transcriptional dynamics | Lotka–Volterra | *E. coli* telegraph |
| --- | --- | --- | --- |
| Observations $N$ | 512 cells | 128 trajectories | 3833 cells |
| Covariates | length $L$, site $z$ | initial counts $(X_0, Y_0)$ | fixation time $t_i$ |
| Inferred parameters | $(k_+, r_{\text{burst}}, D)$ | $(\alpha, \beta, \gamma)$ | $(K_{1,2}, K_{2,1}, B_1, B_2)$ |
| Generating values | $(15, 10, 0.1)$ | $(5, 0.02, 5)$ | real data |
| Fixed parameter | $k_- = 5$ | $\eta = 1$ | $k_- = 0.00533 \text{ s}^{-1}$ |
| Summaries per observation | mRNA count; its squared deviation from the batch mean; s.d. of mRNA positions (3) | log mean, log variance and lag-0.1 autocorrelation of $X$ and of $Y$; cross-correlation (7) | $\mathbf{1}\{\text{count} \leq m\}$, $m = 0, 1, 3, 7, 15$, in the block of the cell’s fixation time (80) |
| Particles $P$ | 500 | 1000 | 1000 |
| Fixed batch sizes | 32, 64, 128, 256, 512 | 8, 16, 32, 64, 128 | 64, 256, 1024, 3833 |
| Adaptive initial batch $n_0$ | 2, 4, 8, 16, 32 | 2, 4, 8, 16, 32 | 16, 64, 256 |
| Smoothing weight $\lambda$ | 0, 0.25, 0.5, 0.75, 1 | 0, 0.25, 0.5, 0.75, 1 | 1 |
| Noise ratio $c$ | 0.1, 0.2, 0.3, 0.4, automatic | 0.1, 0.2, 0.3, 0.4, automatic | automatic |

After computing the perturbation covariance Γ_*t*_ in Algorithm 1, we calculate the effective sample size 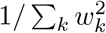. If it is at most *P/*2, we resample *P* particles with replacement according to their weights and reset the weights to 1*/P* before forming *q*_*t*+1_, keeping Γ_*t*_ at its pre-resampling value.

The hyperparameter sweeps (**Figs**. 4 and 9) use one fixed synthetic dataset per model and three independently seeded fits per setting. The shared adaptive setting is *n*_0_ = 2, automatic *c*, and *λ* = 1; each sweep varies one of *n*_0_, *c*, and *λ* over the values in **Tab**. 1. The kinetic and heterogeneity sweeps (**Figs**. 6, S3, S5 and S6) regenerate the dataset for each of ten paired data/SMC seeds per setting. The dataset-size comparison (**Fig**. 5) uses TD datasets of *N* = 512, 2,048, 8,192, and 32,768 cells at the baseline generating values, three seeds each, with the same stopping rule and a 13-hour wall limit. One full-batch fit at *N* = 32,768 reached this limit after passing every target. The targets are *W* = 30, 3, 0.3, and 0.03, and a seed that stopped above a target contributes no point. The *E. coli* comparison uses five runs of each method.

## Supplementary figures

**Fig. S1:**
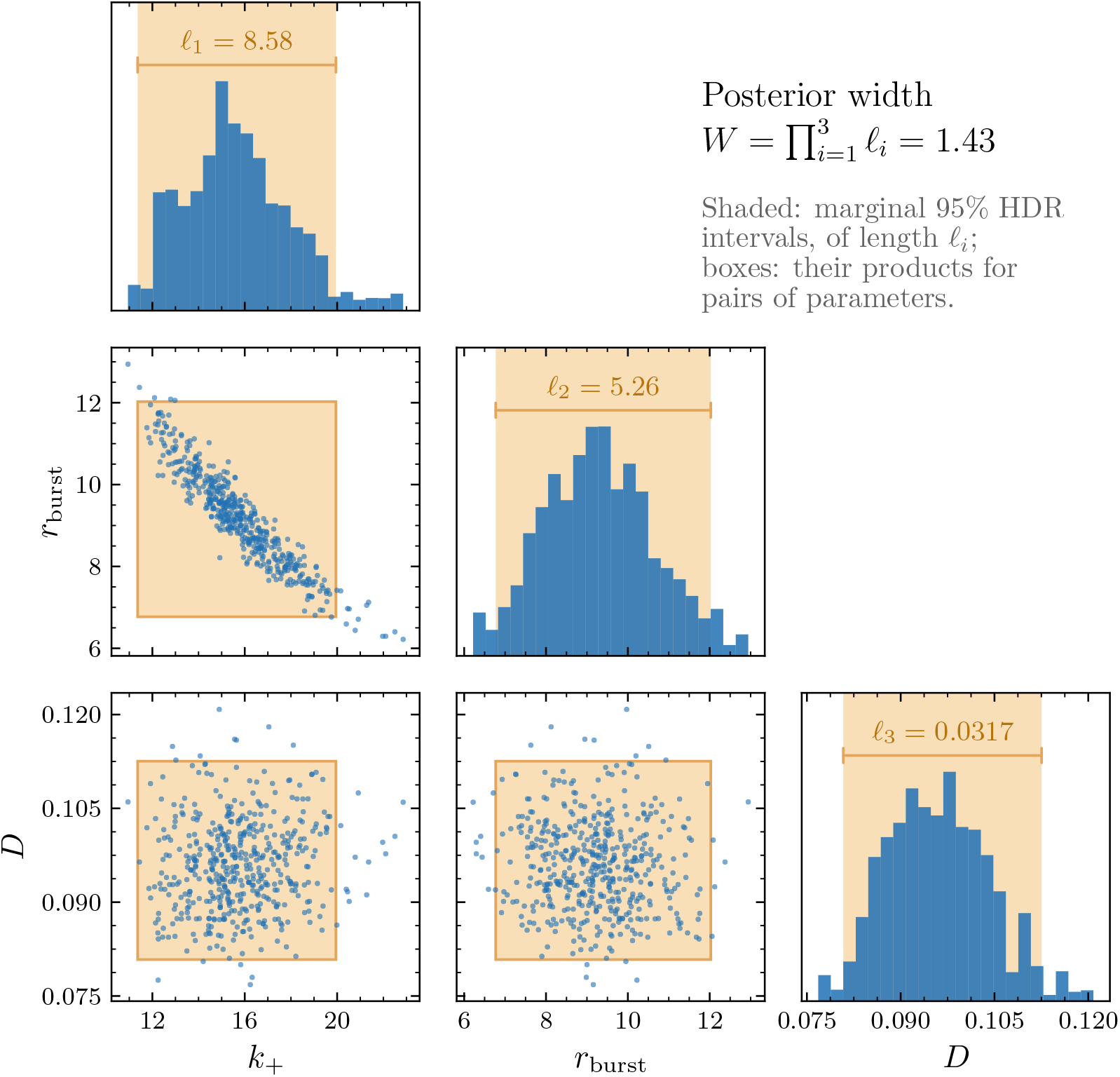
The posterior width *W* . Corner plot of one adaptive posterior for the transcriptional dynamics model (*n*_0_ = 2, automatic *c*, stopping generation of one seed). Shaded intervals are the marginal 95% highest-density intervals, of lengths *l*_*j*_; boxes show their products for pairs of parameters. The reported width is the product of all marginal lengths, *W* = Π_*j*_ *l*_*j*_.

**Fig. S2:**
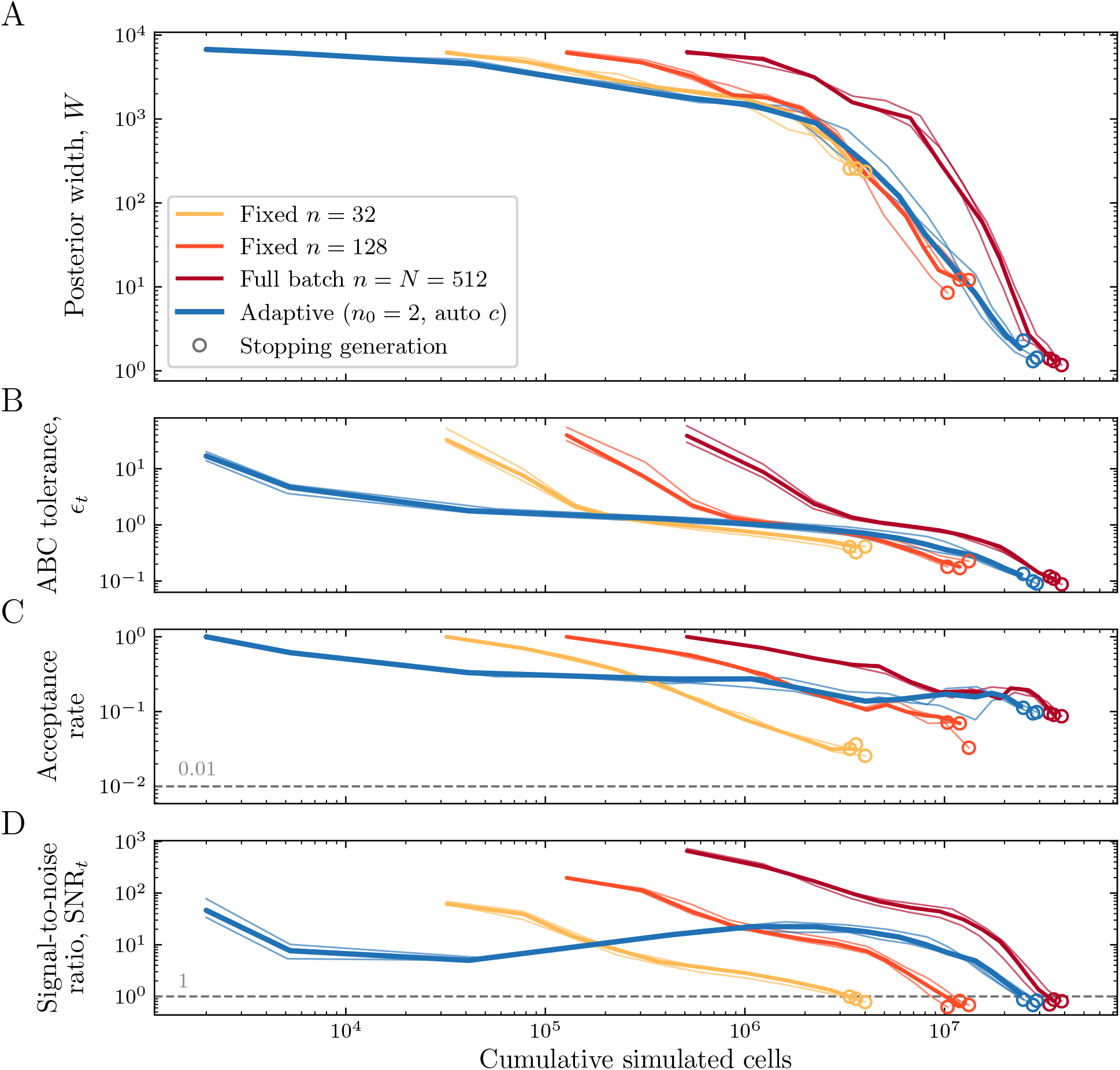
Simulation cost in the transcriptional dynamics model. (A) Posterior width *W* , (B) ABC tolerance, (C) acceptance rate, and (D) signal-to-noise ratio against cumulative simulated cells for the adaptive rule, fixed minibatches, and the full batch on the same dataset. Thin curves show individual seeds and thick curves show medians on common generation support; circles mark each seed’s stopping generation. Dashed lines mark the stopping thresholds. Tolerances are comparable only within a batch size: the distance carries a noise term of order *V/n*, so a small fixed batch reaches a given tolerance with a much broader posterior and stops when its SNR reaches one.

**Fig. S3:**
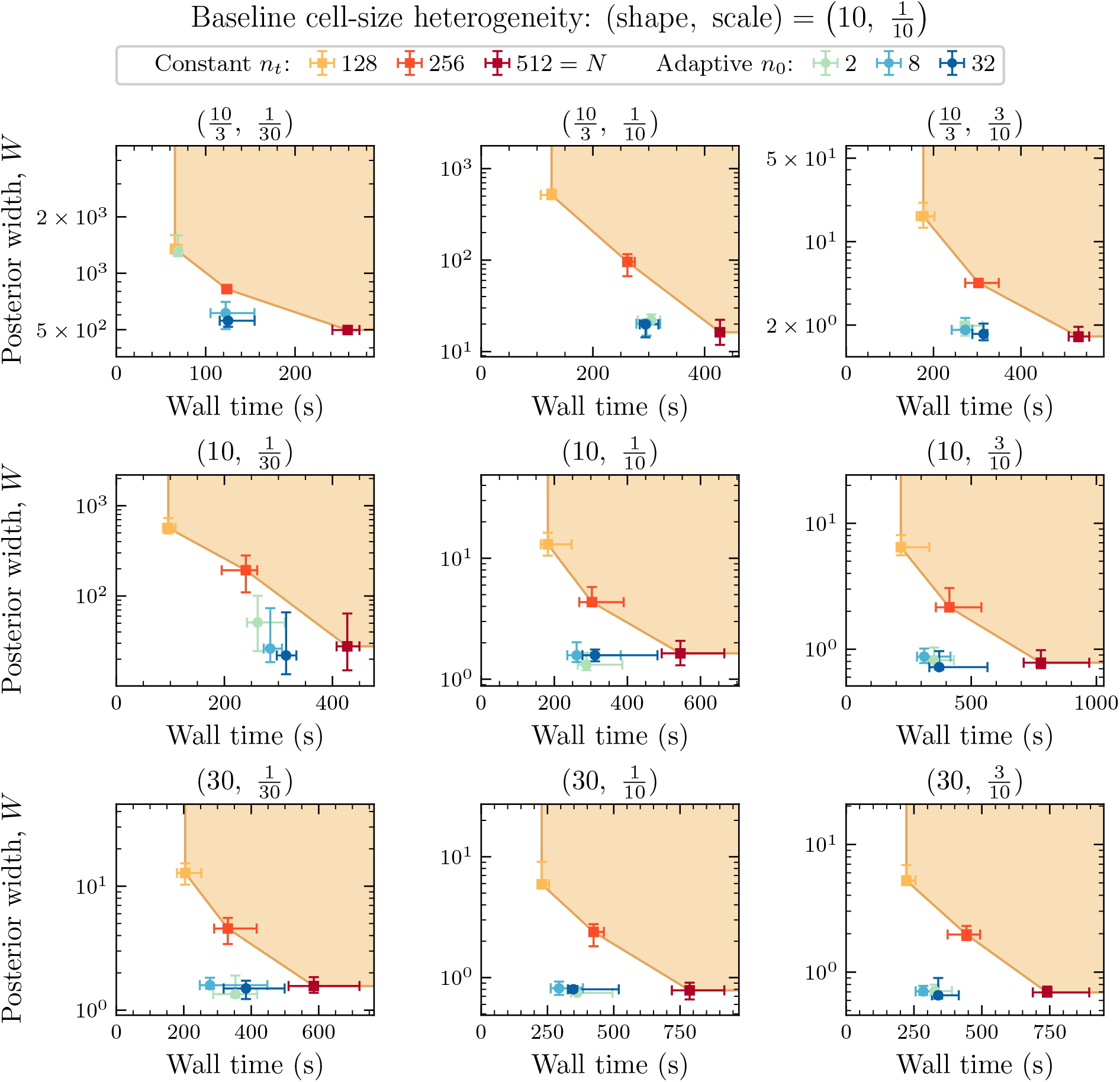
Adaptive and fixed minibatches under changes to the Gamma distribution of nuclear lengths. Each panel shows wall time at the stopping generation against the posterior width *W* for one (shape, scale) pair, as in **Fig**. 6. Points are medians over ten paired data/SMC seeds with quartile bars.

**Fig. S4:**
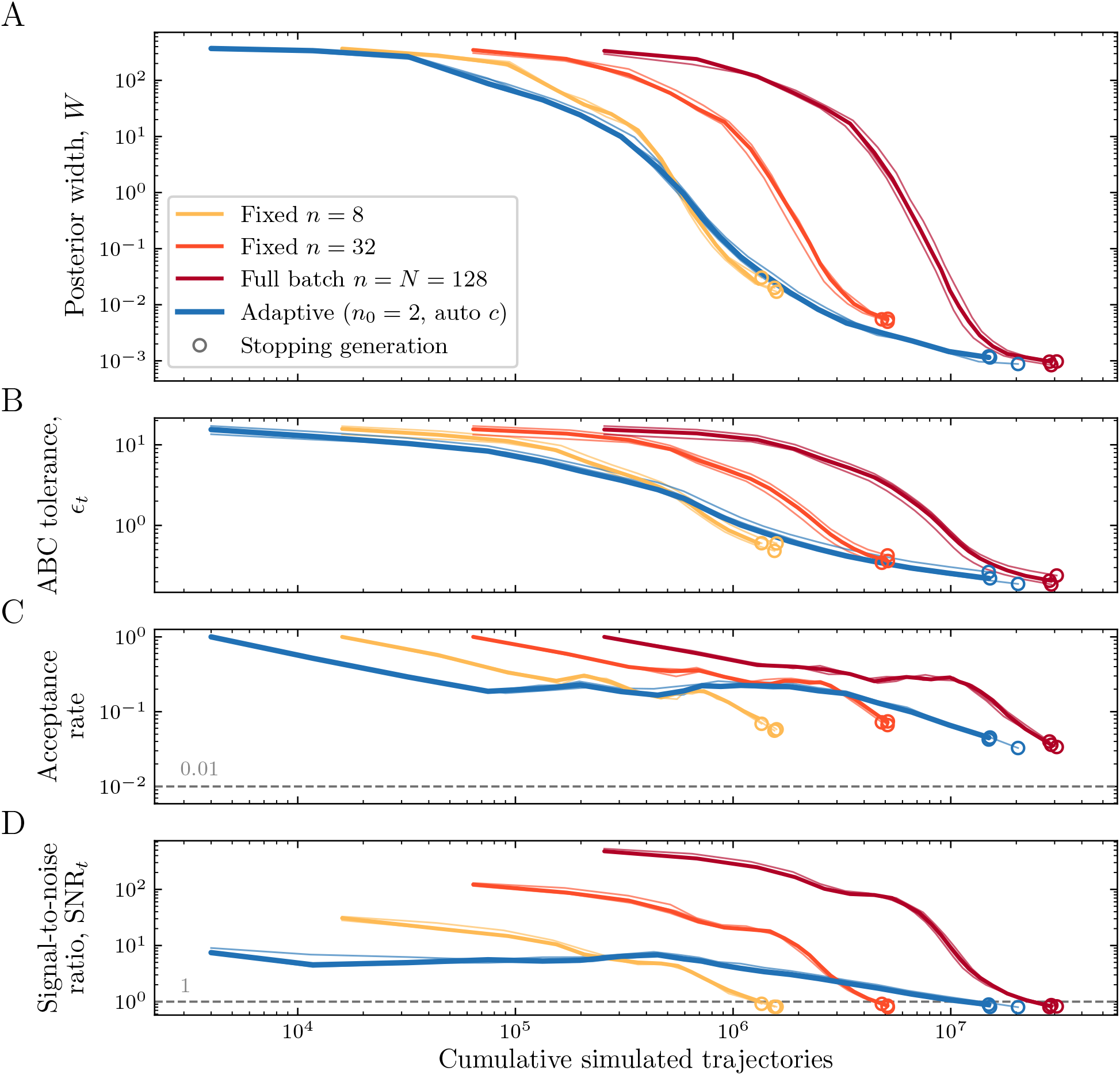
Simulation cost in the stochastic Lotka–Volterra model. (A) Posterior width *W* , (B) ABC tolerance, (C) acceptance rate, and (D) signal-to-noise ratio against cumulative simulated trajectories for the adaptive rule, fixed minibatches, and the full batch on the same dataset, as in **Fig**. S2.

**Fig. S5:**
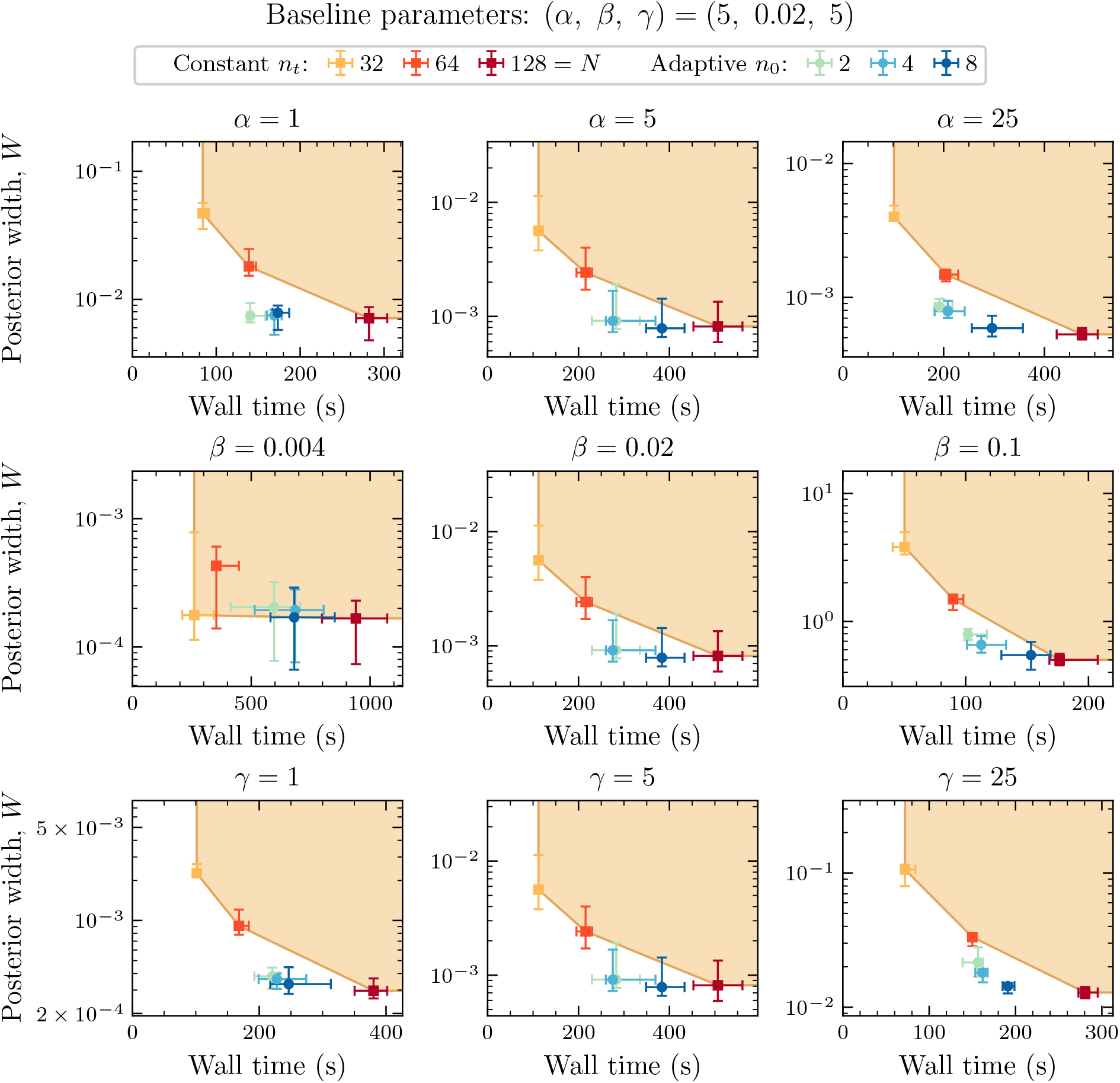
Robustness of the stochastic Lotka–Volterra comparison to changes in kinetic parameters. Each panel varies one of *α, β*, or *γ* with the others at their baseline values and shows wall time at the stopping generation against the posterior width *W* , as in **Fig**. 6. Points are medians over ten paired data/SMC seeds with quartile bars.

**Fig. S6:**
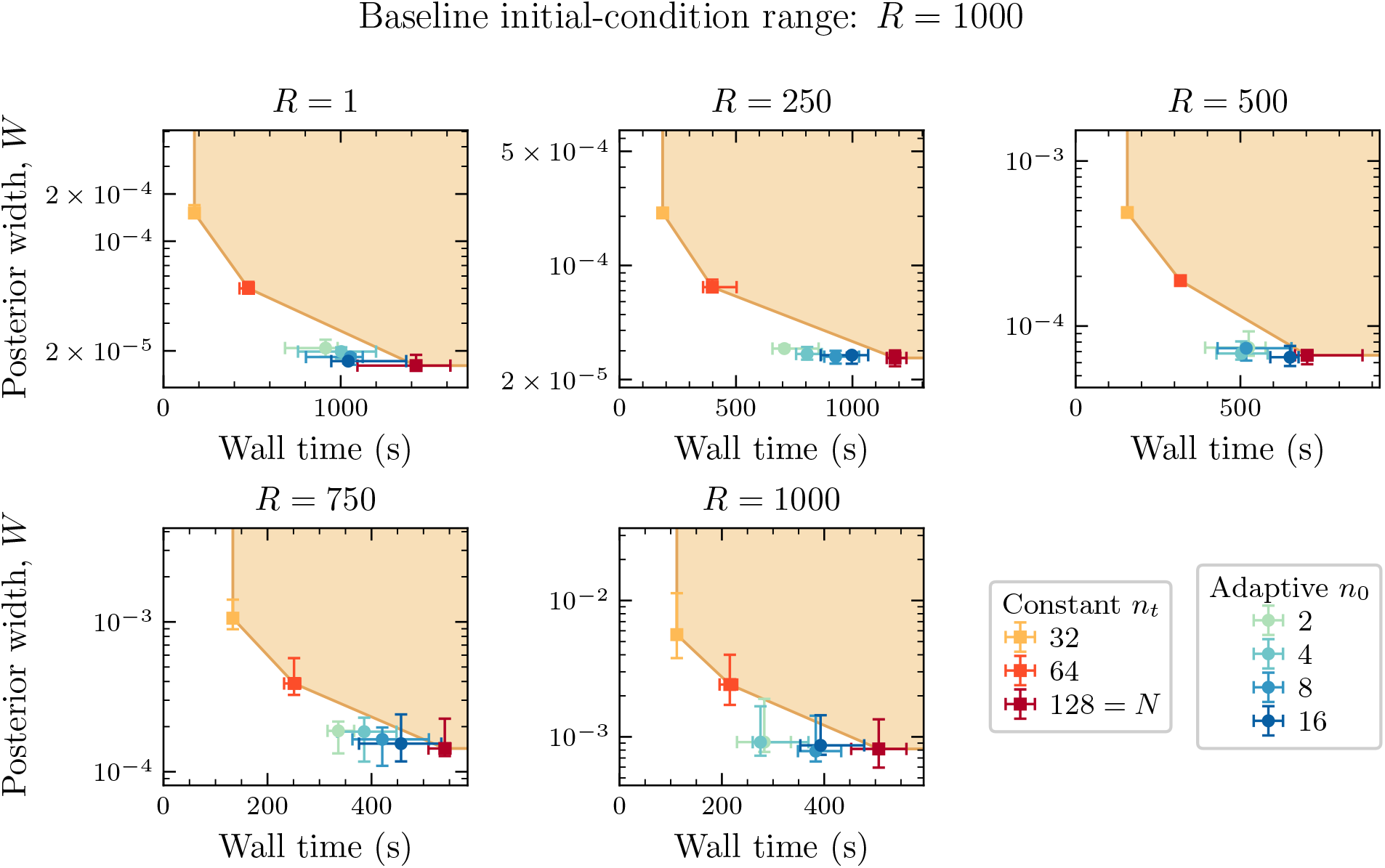
Robustness of the stochastic Lotka–Volterra comparison to the range *R* of initial populations. Each panel shows wall time at the stopping generation against the posterior width *W* for one range, as in **Fig**. 6. Points are medians over ten paired data/SMC seeds with quartile bars.

**Fig. S7:**
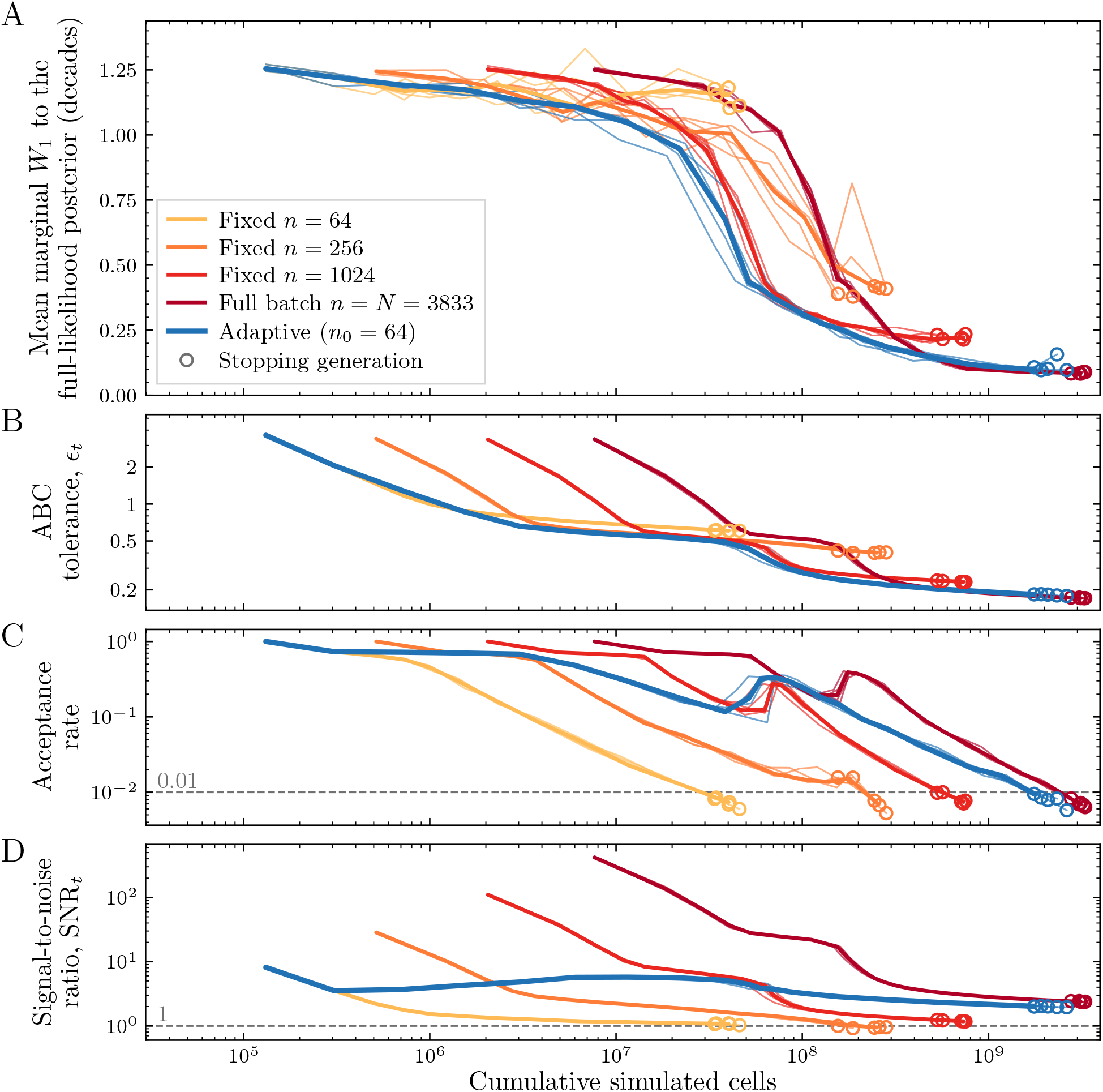
Simulation cost in the *E. coli* application. (A) Mean marginal Wasserstein distance to the likelihood-based posterior, (B) ABC tolerance, (C) acceptance rate, and (D) signal-to-noise ratio against cumulative simulated cells. Thin curves show the five individual runs and thick curves the generation-wise median over the runs that reach each generation, drawn to each method’s median stopping generation, as in **Fig**. S2; circles mark each run’s stopping generation and dashed lines the stopping thresholds.

**Fig. S8:**
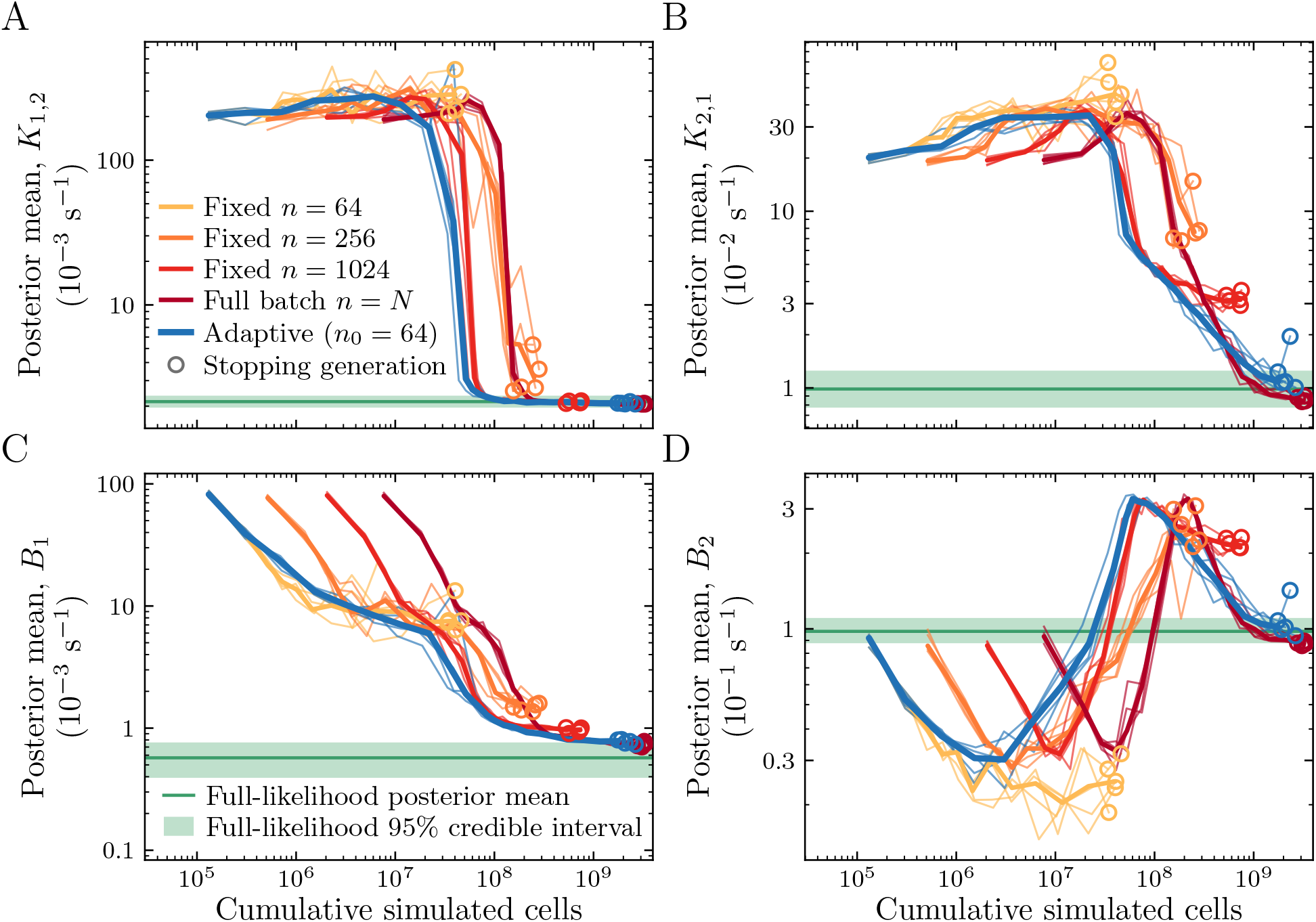
Point estimates against simulation cost in the *E. coli* application. Each panel shows the posterior mean of one rate, (A) *K*_1,2_, (B) *K*_2,1_, (C) *B*_1_, (D) *B*_2_, at every generation against cumulative simulated cells, for the adaptive rule, fixed minibatches, and the full batch. The green line is the mean of the likelihood-based posterior and the band its central 95% credible interval. Rates are on a logarithmic axis, in the units of **Fig**. 10C. Thin curves show the five individual runs and thick curves the generation-wise median, drawn to each method’s median stopping generation; circles mark each run’s stopping generation, as in **Fig**. S7.

## References

1. Jørgensen, A. C. S., Ghosh, A., Sturrock, M. & Shahrezaei, V. Efficient Bayesian inference for stochastic agent-based models. PLOS Computational Biology 18, e1009508 (2022). doi:10.1371/journal.pcbi.1009508.

2. Cranmer, K., Brehmer, J. & Louppe, G. The frontier of simulation-based inference. Proceedings of the National Academy of Sciences 117, 30055–30062 (2020). doi:10.1073/pnas.1912789117.

3. Wang, X., Kelly, R. P., Jenner, A. L., Warne, D. J. & Drovandi, C. A comprehensive guide to simulation-based inference in computational biology. arXiv preprint (2024). doi:10.48550/arXiv.2409.19675.

4. Gough, A., Stern, A. M., Maier, J., Lezon, T., Shun, T.-Y., Chennubhotla, C., Schurdak, M. E., Haney, S. A. & Taylor, D. L. Biologically relevant heterogeneity: metrics and practical insights. SLAS Discovery 22, 213–237 (2017). doi:10.1177/2472555216682725.

5. Zechner, C., Unger, M., Pelet, S., Peter, M. & Koeppl, H. Scalable inference of heterogeneous reaction kinetics from pooled single-cell recordings. Nature Methods 11, 197–202 (2014). doi:10.1038/nmeth.2794.

6. Stallvik, A.Kaltenbach, H.-M. & Stelling, J. Optimal input design for model selection in systems with cell-to-cell variability Computational Methods in Systems Biology (CMSB 2025) 15959 (Springer Nature Switzerland, Cham, 2025), 240–257. doi:10.1007/978-3-032-01436-8_13.

7. Lueckmann, J.-M., Boelts, J., Greenberg, D., Goncalves, P. & Macke, J. Benchmarking simulation-based inference Proceedings of the 24th International Conference on Artificial Intelligence and Statistics 130 (PMLR, 2021), 343–351. https://proceedings.mlr.press/v 130/lueckmann21a.html.

8. Schälte, Y. & Hasenauer, J. Efficient exact inference for dynamical systems with noisy measurements using sequential approximate Bayesian computation. Bioinformatics 36, i551–i559 (2020). doi:10.1093/bioinformatics/btaa397.

9. Alamoudi, E., Schälte, Y., Müller, R., Starruß, J., Bundgaard, N., Graw, F., Brusch, L. & Hasenauer, J. FitMultiCell: simulating and parameterizing computational models of multi-scale and multi-cellular processes. Bioinformatics 39, btad674 (2023). doi:10.1093/bioinformatics/btad674.

10. Osuntoki, I. G., Harrison, A., Dai, H., Bao, Y. & Zabet, N. R. HiCPotts: an R/Bioconductor package to identify significant interactions in chromosome conformation capture data and model sources of bias. Bioinformatics, btag673 (2026). doi:10.1093/bioinformatics/btag673.

11. Li, X., Chadwick, F. & Swallow, B. Advances in approximate Bayesian inference for models in epidemiology. Epidemics 53, 100855 (2025). doi:10.1016/j.epidem.2025.100855.

12. Toni, T., Welch, D., Strelkowa, N., Ipsen, A. & Stumpf, M. P. H. Approximate Bayesian computation scheme for parameter inference and model selection in dynamical systems. Journal of The Royal Society Interface 6, 187–202 (2009). doi:10.1098/rsif.2008.0172.

13. Beaumont, M. A., Cornuet, J.-M.Marin, J.-M. & Robert, C. P. Adaptive approximate Bayesian computation. Biometrika 96, 983–990 (2009). doi:10.1093/biomet/asp052.

14. Del Moral, P., Doucet, A. & Jasra, A. An adaptive sequential Monte Carlo method for approximate Bayesian computation. Statistics and Computing 22, 1009–1020 (2012). doi:10.1007/s11222-011-9271-y.

15. Fearnhead, P. & Prangle, D. Constructing summary statistics for approximate Bayesian computation: semi-automatic approximate Bayesian computation. Journal of the Royal Statistical Society Series B: Statistical Methodology 74, 419–474 (2012). doi:10.1111/j.1467-9868.2011.01010.x.

16. Prangle, D. Adapting the ABC distance function. Bayesian Analysis 12, 289–309 (2017). doi:10.1214/16-BA1002.

17. Everitt, R. G. & Rowińska, P. A. Delayed acceptance ABC-SMC. Journal of Computational and Graphical Statistics 30, 55–66 (2021). doi:10.1080/10618600.2020.1775617.

18. Kerama, I., Thorne, T. & Everitt, R. G. Rare event ABC-SMC2. arXiv preprint (2022). doi:10.48550/arXiv.2211.02172.

19. Prescott, T. P. & Baker, R. E. Multifidelity approximate Bayesian computation with sequential Monte Carlo parameter sampling. SIAM/ASA Journal on Uncertainty Quantification 9, 788– 817 (2021). doi:10.1137/20M1316160.

20. Steele, T. P. & Warne, D. J. Simulation and inference methods for non-Markovian stochastic reaction networks. PLOS Computational Biology 22, e1014687 (2026). doi:10.1371/journal.pcbi.1014687.

21. Bottou, L., Curtis, F. E. & Nocedal, J. Optimization methods for large-scale machine learning. SIAM Review 60, 223–311 (2018). doi:10.1137/16M1080173.

22. Martina Perez, S., Sailem, H. & Baker, R. E. Efficient Bayesian inference for mechanistic modelling with high-throughput data. PLOS Computational Biology 18, e1010191 (2022). doi:10.1371/journal.pcbi.1010191.

23. Pillai, N. S. & Smith, A. Ergodicity of approximate MCMC chains with applications to large data sets. arXiv preprint (2015). doi:10.48550/arXiv.1405.0182.

24. Bardenet, R., Doucet, A. & Holmes, C. On Markov chain Monte Carlo methods for tall data. Journal of Machine Learning Research 18, 1–43 (2017). https://www.jmlr.org/papers/v18/15-205.html.

25. Bardenet, R., Doucet, A. & Holmes, C. Towards scaling up Markov chain Monte Carlo: an adaptive subsampling approach Proceedings of the 31st International Conference on Machine Learning 32 (2014), 405–413. https://proceedings.mlr.press/v32/bardenet14.html.

26. Byrd, R. H., Chin, G. M., Nocedal, J. & Wu, Y. Sample size selection in optimization methods for machine learning. Mathematical Programming 134, 127–155 (2012). doi:10.1007/s10107-012-0572-5.

27. Drovandi, C. C. & Pettitt, A. N. Estimation of parameters for macroparasite population evolution using approximate Bayesian computation. Biometrics 67, 225–233 (2011). doi:10.1111/j.1541-0420.2010.01410.x.

28. Wilkinson, R. D. Approximate Bayesian computation (ABC) gives exact results under the assumption of model error. Statistical Applications in Genetics and Molecular Biology 12, 129–141 (2013). doi:10.1515/sagmb-2013-0010.

29. Gorin, G., Vastola, J. J. & Pachter, L. Studying stochastic systems biology of the cell with single-cell genomics data. Cell Systems 14, 822–843.e22 (2023). doi:10.1016/j.cels.2023.08.004.

30. Miles, C. E. Incorporating spatial diffusion into models of bursty stochastic transcription. Journal of the Royal Society Interface 22, 20240739 (2025). doi:10.1098/rsif.2024.0739.

31. Miles, C. E., McKinley, S. A., Ding, F. & Lehoucq, R. B. Inferring stochastic rates from heterogeneous snapshots of particle positions. Bulletin of Mathematical Biology 86, 74 (2024). doi:10.1007/s11538-024-01301-4.

32. Raj, A., van den Bogaard, P., Rifkin, S. A., van Oudenaarden, A. & Tyagi, S. Imaging individual mRNA molecules using multiple singly labeled probes. Nature Methods 5, 877–879 (2008). doi:10.1038/nmeth.1253.

33. Gillespie, D. T. Approximate accelerated stochastic simulation of chemically reacting systems. The Journal of Chemical Physics 115, 1716–1733 (2001). doi:10.1063/1.1378322.

34. Wang, M., Zhang, J., Xu, H. & Golding, I. Measuring transcription at a single gene copy reveals hidden drivers of bacterial individuality. Nature Microbiology 4, 2118–2127 (2019). doi:10.1038/s41564-019-0553-z.

35. Kilic, Z., Schweiger, M., Moyer, C., Shepherd, D. & Pressé, S. Gene expression model inference from snapshot RNA data using Bayesian non-parametrics. Nature Computational Science 3, 174–183 (2023). doi:10.1038/s43588-022-00392-0.

36. Peccoud, J. & Ycart, B. Markovian modeling of gene-product synthesis. Theoretical Population Biology 48, 222–234 (1995). doi:10.1006/tpbi.1995.1027.

37. Gillespie, D. T. Exact stochastic simulation of coupled chemical reactions. The Journal of Physical Chemistry 81, 2340–2361 (1977). doi:10.1021/j100540a008.

38. Munsky, B. & Khammash, M. The finite state projection algorithm for the solution of the chemical master equation. The Journal of Chemical Physics 124, 044104 (2006). doi:10.1063/1.2145882.

39. Warne, D. J., Zhu, X., Steele, T. P., Johnston, S. T., Sisson, S. A., Faria, M., Murphy, R. J. & Browning, A. P. A multilevel hierarchical framework for quantification of experimental heterogeneity in population snapshot data. PLOS Computational Biology 22, e1014379 (2026). doi:10.1371/journal.pcbi.1014379.

40. Warne, D. J., Prescott, T. P., Baker, R. E. & Simpson, M. J. Multifidelity multilevel Monte Carlo to accelerate approximate Bayesian parameter inference for partially observed stochastic processes. Journal of Computational Physics 469, 111543 (2022). doi:10.1016/j.jcp.2022.111543.

41. Papamakarios, G. & Murray, I. Fast ε-free inference of simulation models with Bayesian conditional density estimation Advances in Neural Information Processing Systems 29 (2016). https://papers.nips.cc/paper_files/paper/2016/hash/6aca97005c68f1206823815f66102863-Abstract.html.

42. Greenberg, D., Nonnenmacher, M. & Macke, J. Automatic posterior transformation for likelihood-free inference Proceedings of the 36th International Conference on Machine Learning 97 (2019), 2404–2414. https://proceedings.mlr.press/v97/greenberg19a.html.

43. Zhang, R., Chkrebtii, O. A. & Xiu, D. Likelihood-free posterior density learning for uncertainty quantification in inference problems. arXiv preprint (2025). doi:10.48550/arXiv.2508.00167.

44. Haario, H., Saksman, E. & Tamminen, J. An adaptive Metropolis algorithm. Bernoulli 7, 223–242 (2001). doi:10.2307/3318737.

